# *De novo* genome assembly and transcriptome analysis for the drought and salt resistant *Solanum sitiens*

**DOI:** 10.1101/2020.06.14.151043

**Authors:** C. Molitor, T.J. Kurowski, P.M. Fidalgo de Almeida, P. Eerolla, D.J. Spindlow, S.P. Kashyap, B. Singh, H.C. Prasanna, A.J. Thompson, F.R. Mohareb

## Abstract

*Solanum sitiens* is a self-incompatible wild relative of tomato, characterised by salt and drought resistance traits, with the potential to contribute to crop improvement in cultivated tomato. This species has a distinct morphology, classification and ecotype compared to other stress resistant wild tomato relatives such as *S. pennellii* and *S. chilense*. Therefore, the availability of a high-quality reference genome for *S. sitiens* will facilitate the genetic and molecular understanding of salt and drought resistance. Here, we present a *de novo* genome and transcriptome assembly for *S. sitiens* (Accession LA1974). A hybrid assembly strategy was followed using Illumina short reads (∼159X coverage) and PacBio long reads (∼44X coverage), generating a total of ∼262 Gbp of DNA sequence; in addition, ∼2,670 Gbp of BioNano data was obtained. A reference genome of 1,245 Mbp, arranged in 1,481 scaffolds with a N50 of 1,826 Mbp was generated. Genome completeness was estimated at 95% using the Benchmarking Universal Single-Copy Orthologs (BUSCO) and the K-mer Analysis Tool (KAT); this is within the range of current high-quality reference genomes for other tomato wild relatives. Additionally, we identified three large inversions compared to *S. lycopersicum*, containing several drought resistance related genes, such as *beta-amylase 1* and *YUCCA7*.

In addition, ∼63 Gbp of RNA-Seq were generated to support the prediction of 31,164 genes from the assembly, and perform a *de novo* transcriptome. Some of the protein clusters unique to *S. sitiens* were associated with genes involved in drought and salt resistance, including *GLO1* and *FQR1*.

This first reference genome for *S. sitiens* will provide a valuable resource to progress QTL studies to the gene level, and will assist molecular breeding to improve crop production in water-limited environments.

## Introduction

Cultivated tomato (*S. lycopersicum* L.) is grown commercially as an irrigated field crop for processing and fresh fruit production. Relative to many crops it is sensitive to soil water deficits, especially at flowering [1] and pressure on water resources [2] make high yield under water deficit conditions, and high water use efficiency (yield per unit water input) important breeding targets [3] [4]. The physiological traits required to achieve these broad targets are complex [5] and require deep understanding of water flows within the soil-plant-atmosphere system and phenology (relationship between crop development and climate); smart breeding requires a knowledge of the genetic loci and their interactions to control specific contributing plant traits.

Crop wild relatives are an important source of genetic variation for crop improvement; they have been deployed successfully to improve e.g. disease resistance, flavour and fruit size in cultivated tomato in recent decades [6]; there remains the opportunity to transfer adaptations to abiotic stresses such as drought and salinity from wild species to tomato crops. Tomato and its closest wild relatives are classified into *Solanum* section *Lycopersicon* containing 13 species [7], including three species with drought resistant accessions: *S. pennellii, S. pimpinellifolium* and *S. chilense* [3] [8]. There is a fourth drought resistant species, *S. sitiens* [9], but this is considered to be within an outgroup, *Solanum* sect. *Lycopersicoides*. The latter contains only one other species, *S. lycopersicoides* [7], and both members of this section can be hybridised with cultivated tomato by overcoming significant reproductive barriers [10] [11] [12].

Of the drought resistant species, excellent genetic and genomic resources (including *de novo* assembly) exist for *S. pennellii* [13] [14], where highly successful strategies have been employed to exploit allelic variation [15], and assemblies have been recently reported for *S. chilense* LA3111 [16] and *S. pimpinellifolium* LA0480 [17]. Extensive resequencing data mapped against cultivated tomato is available from over 500 accessions, including all wild species [18] [19] [20], and a pan-genome has been created for tomato using 725 accessions [21], but this included limited numbers of wild species and only within the *Solanum* sect. *Lycopersicon*. A genome assembly for *S. sitiens* is lacking, therefore, the aim of this work was to create this genomic resource to underpin use of this species in genetic studies of drought and salinity resistance.

*Solanum sitiens* is found in an extremely dry habitat of Chile, within a very limited geographic range on the plateau of the hyper arid Atacama Desert, with most accessions collected between 2,500 and 3,300 m in altitude [9] [22]. The C.M. Rick Tomato Genetic Resources Centre (University of California at Davis), indicates resistance to drought stress as a general feature of *S. sitiens*, and recommends accessions LA1974 and LA2876 for investigations related to drought. Accession LA1974, the object of this study, was collected by Carlos Ochoa from Chuquicamata, Antofagasta, Chile in 1979 from an extremely dry habitat. The collection sites recorded for *S. sitiens* are typically arid and with soil salinity at levels where cultivated tomatoes would not be productive [9]. Descriptions of the morphological adaptations of *S. sitiens* to drought are reported: thick leathery leaves that are small and able to fold along the mid-vein, and the ability to regenerate from the base of the stem after prolonged drought. However, there are no known studies of drought physiology in this species. In addition, *S. sitiens* has a seed dispersal strategy unique within the *Solanum* genus in which fruits desiccate, rather than soften and ripen, and then drop to the ground for dispersal by wind or by rolling down slopes [9].

Recently, a library of 56 introgression lines representing 93% of the genome of *S. sitiens* accessions LA4331 and LA1974 was reported [23]. The availability of a reference genome assembly, in addition to these new genetic resources in *S. sitiens*, will benefit breeding efforts in cultivated tomato and open new opportunities for further studies linking genes to the unique biology of this species. *S. sitiens* is allogamous and self-incompatible, so likely to be highly heterozygous, making genome assembly more challenging than for inbred species.

## Methods

### Plant material

Seeds of *S. sitiens* accession LA1974 were obtained from the C.M. Rick Tomato Genetics Resource Center maintained by the Department of Plant Sciences, University of California, Davis, USA. Seeds were treated with 50% v/v household bleach, equivalent to 2.25% w/v sodium hypochlorite for 60 mins, rinsed in tap water and then germinated on filter paper soaked in distilled water at 25°C in the dark. A single plant was clonally propagated by rooting of shoot cuttings and all sequence data was obtained from this single clone.

### DNA and RNA extraction

Genomic DNA and total RNA for Illumina sequencing was prepared using the DNeasy and RNeasy Plant Mini Kits, respectively (Qiagen, Manchester, UK), according to the manufacturer’s instructions. High molecular weight genomic DNA for PacBio sequencing and for Bionano optical mapping was prepared by the Earlham Institure (previously known as The Genome Analysis Centre, Norwich, UK) using a Bionano Prep Plant Tissue DNA isolation kit, according to manufacturer’s instructions; this involved purification of nuclei which were then embedded in agarose and digested with proteinase K and RNase A before recovery of DNA with agarase.

### Sequencing data

Two PCR-free Paired-End (PE) libraries with a read length of 250 base pairs (bp) and an insert size of 395 bp were prepared for sequencing on an Illumina Hiseq2500™platform at the Earlham Institute using the Whole Genome Sequencing (WGS) approach. The sequencing yielded a total of ∼172 Gbp and the quality of the Illumina reads was assessed with FastQC v0.11 [24].

Long reads were sequenced on two different Pacific Bioscience platforms, RS-II and Sequel. ***RS-II***: 18 Single Molecule, Real-Time (SMRT) cells were sequenced on a PacBio RS-II platform with P6-C4 chemistry. In total, ∼8.7 Gbp of raw sequencing data were generated in ∼1.3 million reads. The N50 of the reads was 10,991 bp. ***Sequel:*** 12 SMRT cells were sequenced on a Sequel Platform. In total, ∼42.4 Gbp of raw sequencing data were generated in ∼4.9 million reads. The reads N50 was 14,244 bp. The outputs from the two platforms were converted to fasta files and merged together for the subsequent analyses. The length distribution of the PacBio reads are available as Supplementary Figure 1.

**Figure 1:**
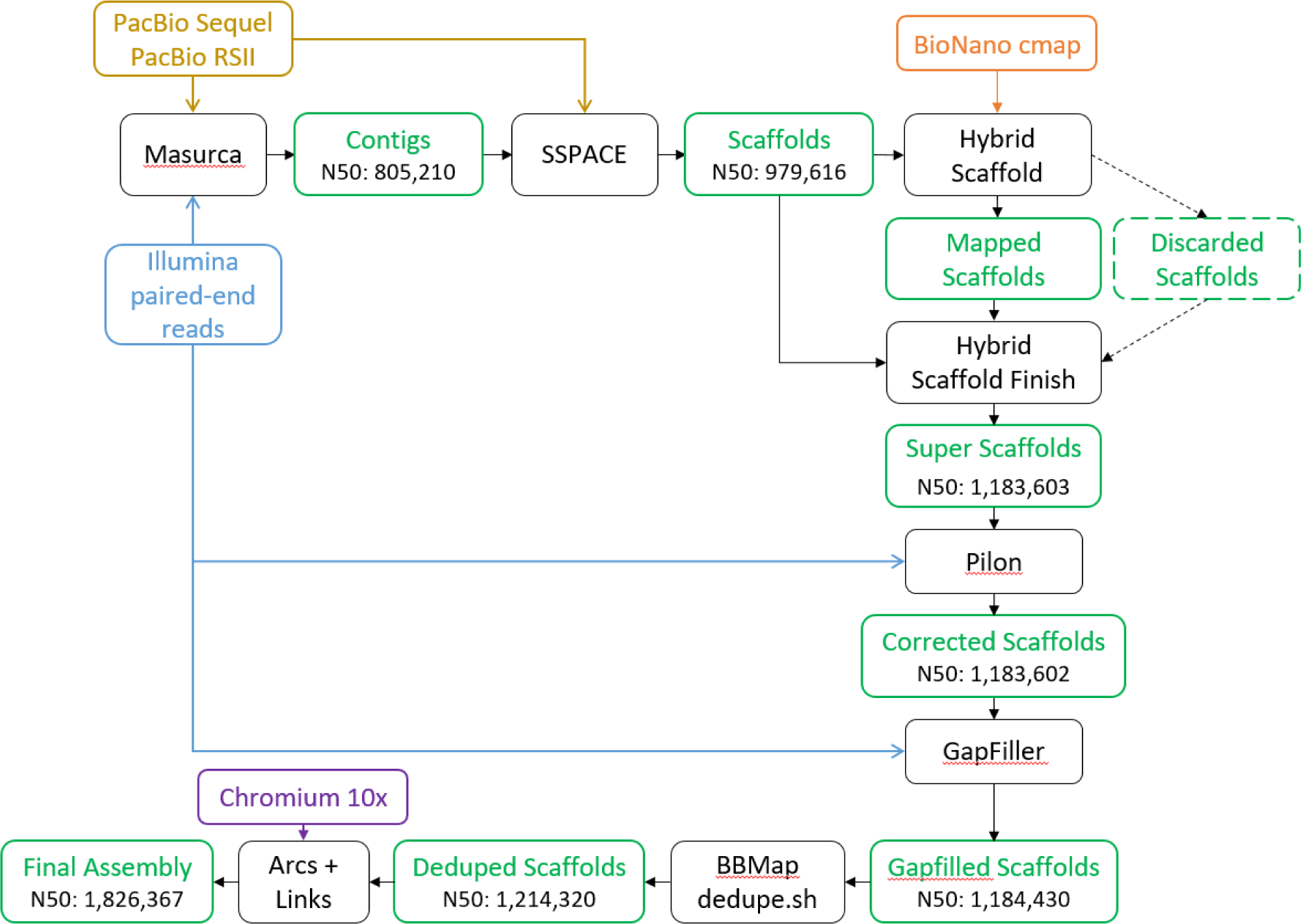
Overview of the assembly pipeline. Tools are in black, assembly steps in green.

Additionally, ∼1,198 Gbp of optical maps longer than 100 kbp were generated on a BioNano Irys platform at the Earlham Institute using the BssS1 nicking enzyme. The raw bnx file was converted to cmap format and produced 4,426 consensus maps [25]. To assist in overcoming the heterozygosity challenge for *S. sitiens* and to generate phased haplotypes reference genome, a single library of PE 10X Chromium data was sequenced from a fresh leaf at the Earlham Institute following 10x Genomics guidelines for genomes between 0.1 and 1.6 Gbp. The resulting fastq files contained ∼40 Gbp of data. The *“basic”* pipeline from LongRanger v2.2.2 interleaved the two fastq files and performed read trimming, barcode error correction and barcode whitelisting. LongRanger also moved the 10x molecule barcode, present in the first 16 bp of each left read, to the corresponding pair read names, a necessary step for most downstream analyses. A custom Perl command (Supplementary Data, “10x_custom_script.pl”) added the barcode to the read name to accommodate Arcs [26] requirements. The reads without a barcode were removed, this filtered ∼13 million reads, corresponding to 5% of the total amount.

12 Illumina paired-end RNA-Seq libraries, generating a total of 63 Gbp of reads were sequenced, first to develop a *de novo* transcriptome assembly and second to generate hints for guided gene prediction from the genome assembly. The quality of the RNA-Seq reads was assessed with FastQC. Pair of reads with at least one read containing unfixable errors, as detected by Rcorrector [27], were removed from the assembly. Rcorrector utilises a k-mer spectra-based methodology to convert rare k-mers within the dataset to trusted k-mers which are more commonly observed within the reads. Here 23-mers were used. Rare k-mers are likely to represent sequencing errors and, although in some cases may be biologically real, were removed to prevent any adverse impact on assembly quality. Trimming of the adapters and low-quality ends from the reads, with a score lower than 5, was done with TrimGalore v0.6.0 [28].

Detailed information on the sequencing throughput and statistics can be found in Supplementary Tables 2-6.

### Genome size estimation

The genome size was estimated with a k-mer based approach [29], Jellyfish v2.2.3 [30], with the “-C” option to consider both strands, counted the k-mers (k=25) present in the two Illumina libraries. The 172 billion bases from the reads generated 129 billion 25-mers. They were plotted as a histogram (see Supplementary Figure 2) which showed two peaks: one at 49 coverage the other at 99, corresponding to the heterozygous and homozygous peaks respectively. The 18.6 billion low coverage 25-mers, below 24 coverage, were considered artefacts and not used as input for the genome size estimation. The remaining 25-mers divided by the homozygous coverage of 99 revealed an estimated genome size of 1,129 Mbp and a heterozygosity of 1.2%. Detailed statistics are available in Supplementary Table 7.

**Figure 2:**
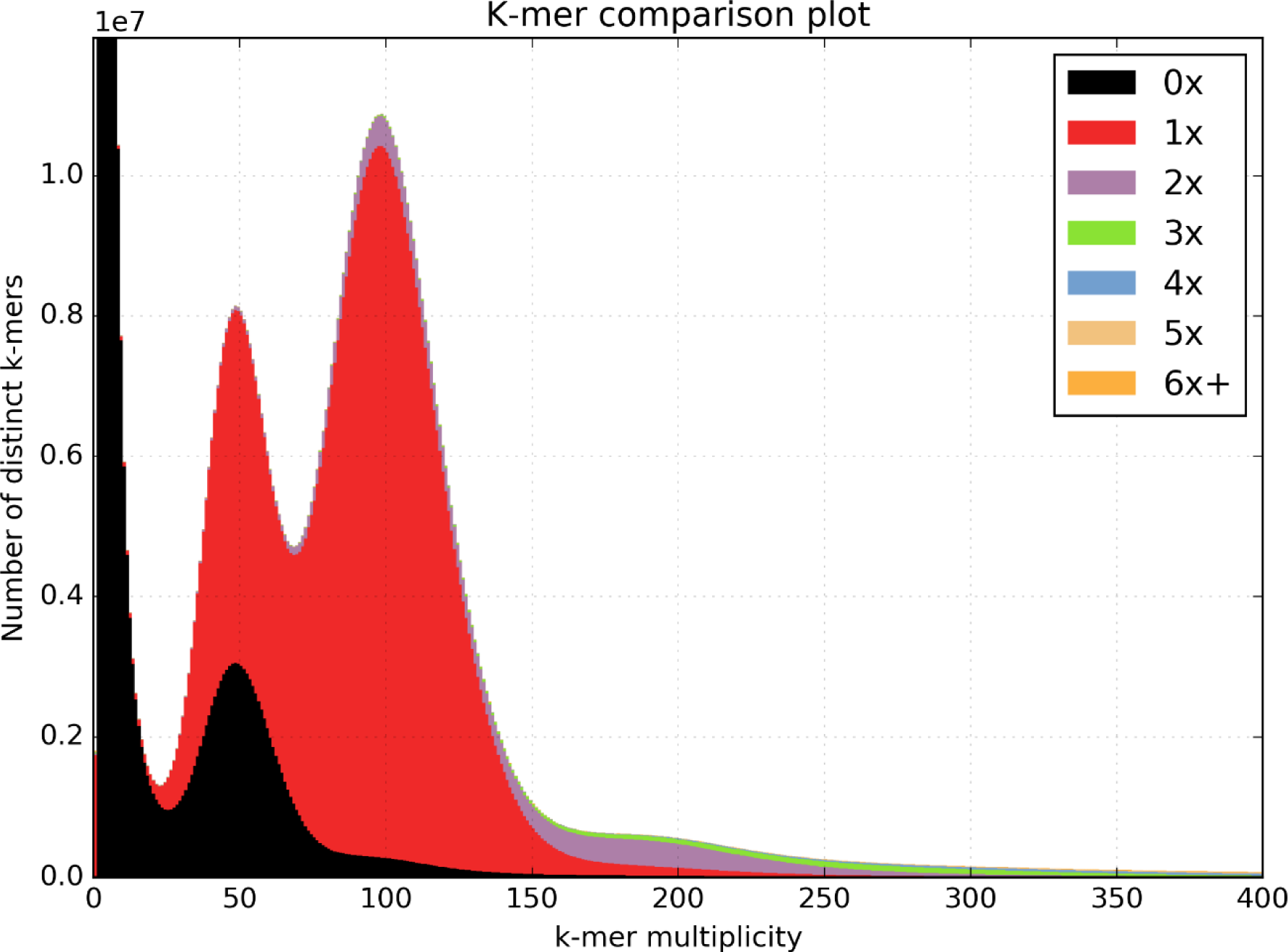
Comparison of the 27mers found in the Illumina libraries, against those from the assembly. The x-axis represents the number of time a specific 27mer has been found in the Illumina reads and the frequency for each occurrence is plotted on the y-axis. The colours represent the copy number of each 27mer in the assembly. The black peak below multiplicity 25 are artefacts, corresponding to 27mers with an error and, as expected, not present in the assembly. As the assembly is producing a single haplotype, the heterozygous and homozygous peaks, at multiplicity 50 and 100 respectively, contains 27mer with a copy number of 1 in the assembly. Similarly, the black peak covering half the heterozygous peak, correspond to the filtered haplotype. The purple peak at multiplicity 200 correspond to repeats present twice in both the reads and the assembly.

### *De novo* genome assembly

The contig assembly was generated with MaSuRCA v3.2.2 [31], a hybrid assembler, taking as input both the Illumina and PacBio reads. Only the PacBio reads longer than 1 kbp were used, the Illumina reads were not trimmed, as advised in MaSuRCA’s documentation. For the deBruijn graph, the optimal k-mer size of 127 bp was automatically determined by MaSuRCA. The k-mer count threshold option was set to 2 as the Illumina coverage was expected to be more than 100x, the number of threads and the jellyfish hash size were set to 64 and 100 billion respectively. The default values were kept for the other parameters. The config file is available as Supplementary Data “masurca_config.txt”.

The contigs were scaffolded with SSPACE v1-1 [32] using 40 threads, the default options and the PacBio reads longer than 1 kbp.

The scaffolds were super-scaffolded with “Hybrid Scaffold” from BioNano Genomics using information from the optical map. The conflict filter levels were set to “no filter” (-B 1 and -N 1), and the configuration xml file (-c) is available as Supplementary Data “hybridScaffold_config.xml”. The cutting enzyme from this project, “BssSI” (CACGAG), was not supported by Hybrid Scaffold and was manually added to the list of available enzymes. The Hybrid Scaffold algorithm discarded all the unmapped scaffolds, damaging the assembly completeness and quality. Hence, the Perl script “hybridScaffold_finish_fasta.pl” script [33] was used to reintegrate them into the assembly.

The polishing of the assembly with the Illumina reads was performed with Pilon v1.22 [34]. Both Illumina libraries were aligned to the scaffolds with the Burrow-Wheeler Aligner (bwa) v0.7.15 using the mem algorithm with the default parameters. Some memory issues were met when running Pilon against the whole assembly, hence we developed a custom script to perform the polishing of the assembly by batches of sequences (available here: https://github.com/MCorentin/run_pilon_batches.sh).

The gaps were filled with GapFiller v1-10 [35], both Illumina libraries were used and defined as having the Forward Reverse (FR) orientation and an insert size of 395 bp, with an standard deviation of 0.25, tolerating insert sizes between 296 and 493 bp. GapFiller was run on 20 threads with 10 iterations and the minimum number of overlapping bases with the edge of the gap was set to 20, the default values were kept for the other parameters.

Some duplications, due to the heterozygous nature of the sample, remained, as shown by the BUSCO results on the assembly (Supplementary Table 1). To remove duplicated scaffolds, the *dedupe*.*sh* script from BBmap v37.32 was run on 60 threads with the following parameters: the *storequality* option was set to false, the *absorbrc* and *touppercase* options were set to true. The scaffolds were considered duplicates if their identity was higher than 90% and they had a minimum overlap of 1000 bases. The allowance for substitutions and edits were determined empirically between a range of 1,000 and 60,000 and 500 and 7,000 respectively. The final values of 40,000 and 5,000 were chosen by assessment of the results with Quast, BUSCO and KAT, see Supplementary Table 9 for more details.

**Table 1:**
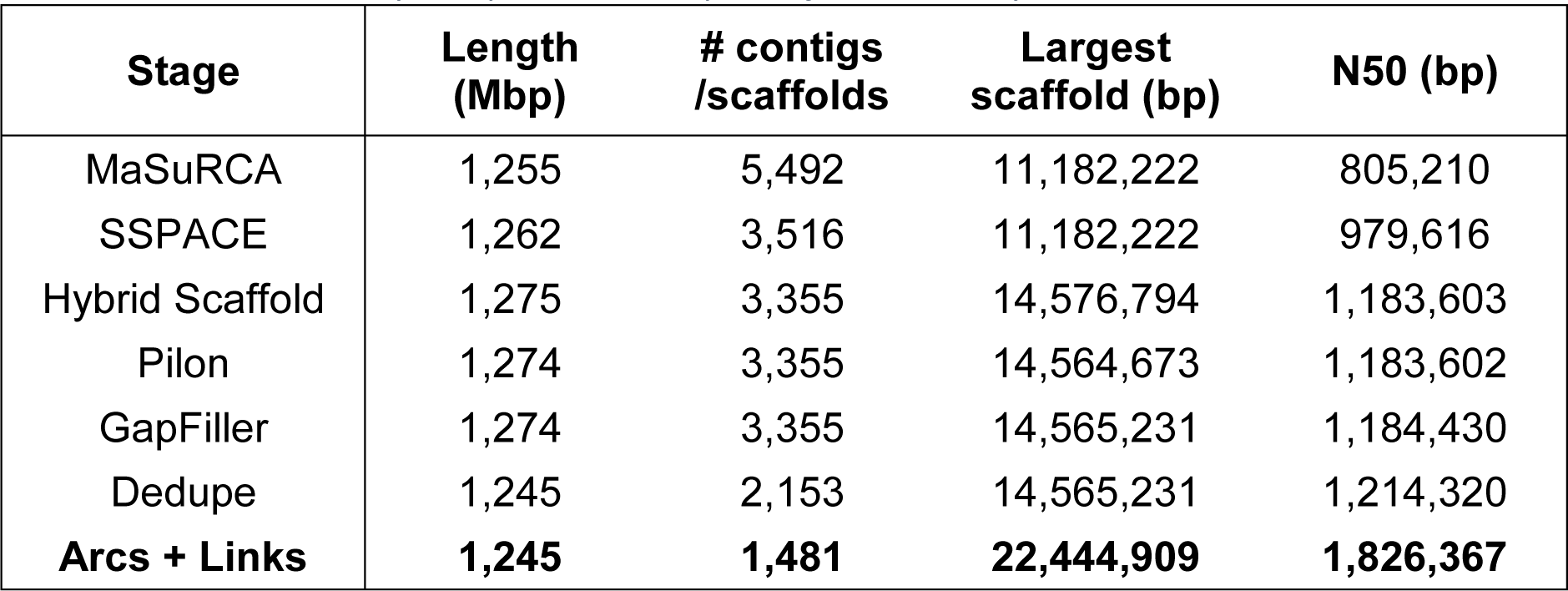
Statistics of the assembly at different stages. Results obtained with Quast v4.5. The length of the assembly is expressed in Mbp. The final assembly statistics are in bold.

| Stage | Length (Mbp) | # contigs /scaffolds | Largest scaffold (bp) | N50 (bp) |
| --- | --- | --- | --- | --- |
| MaSuRCA | 1,255 | 5,492 | 11,182,222 | 805,210 |
| SSPACE | 1,262 | 3,516 | 11,182,222 | 979,616 |
| Hybrid Scaffold | 1,275 | 3,355 | 14,576,794 | 1,183,603 |
| Pilon | 1,274 | 3,355 | 14,564,673 | 1,183,602 |
| GapFiller | 1,274 | 3,355 | 14,565,231 | 1,184,430 |
| Dedupe | 1,245 | 2,153 | 14,565,231 | 1,214,320 |
| **Arcs + Links** | **1,245** | **1,481** | **22,444,909** | **1,826,367** |

**Table 2:**
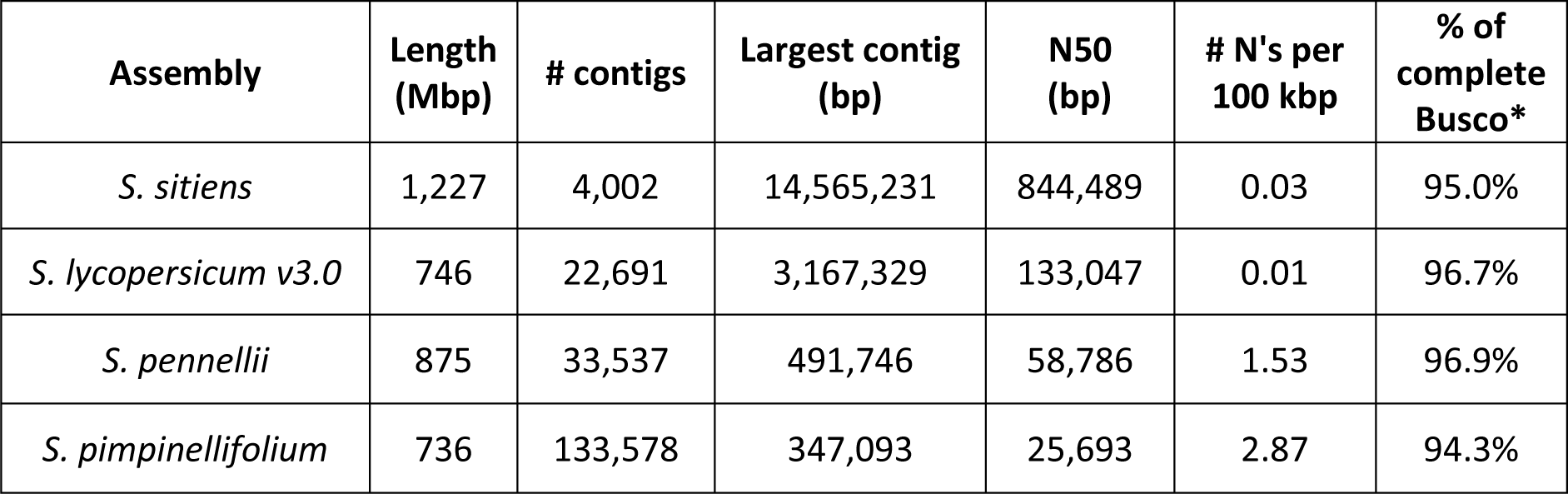
Comparison at the contig level of our S. sitiens assembly against S. pennellii and S. lycopersicum. The scaffolds were broken at every 10 continuous N’s. *The Busco analysis was performed on the original fasta files.

The 10x Genomics Chromium data was used as a final step to super scaffold the assembly with the Arcs and Links [36] pipeline (available at https://github.com/bcgsc/arcs). The Chromium data was aligned to the deduped assembly with bwa v0.7.15 using the mem algorithm with the smart pairing option enabled (-p) to indicate the interleaved nature of the data and otherwise default parameters. First, Arcs v8.25, with default values and harnessing the long-rage information contained in the Chromium data, generated a graph with the scaffolds as nodes and evidence of links between the scaffolds as edges. This graph was transformed to a tsv file with the “makeTSVfile.py” script distributed with Arcs and given as input to Links v1.8.5 for the super scaffolding.

All the commands and scripts necessary to generate the assembly are available at the following github repository, https://github.com/MCorentin/Solanum_sitiens_assembly.

### Assessment of the assembly quality

At each step of the assembly process, results were assessed with Quast v4.5 [37] to measure the sequences length and contiguity, and BUSCO v3.0.2 [38] to estimate the completeness in term of gene content. For BUSCO, the “genome” mode was used to compare the assembly against the 3,052 benchmark genes from the OrthoDB v10 “Solanaceae” dataset [39]. To further confirm the final assembly completeness and duplication rate, the k-mer Analysis Toolkit (KAT) v2.4.0 [40] compared the 27mers presents in the two Illumina libraries against the 27mers from the assembly. Moreover, the RNA-seq data was mapped with STAR v2.6.0c [41] to the draft assembly to validate the correctness and completeness with an independent dataset.

Contamination in the assembly was assessed by aligning the scaffolds, obtained after the GapFiller step, to NCBI’s Non-redundant Protein (NCBI-nr) database with Blastx-fast v2.3.0 [42] with a word size of 20 and otherwise default parameters. The draft assembly was also aligned with Mummer v4.0 [43] to the *S. Lycopersicum* reference genome v3.0 and *S. pennellii*, two closely related species. The nucmer algorithm was used, keeping only the unique anchors in both the reference and query (--mum) and the minimum length of a cluster of matches (-c) was set to 300. The resulting alignment files were filtered with delta-filter only keeping 1-to-1 alignments (-l), to avoid cluttering the dot plots, which were made with mummerplot, using the -large and -layout options.

### Functional annotation and gene characterisation

For the *de novo* transcriptome assembly, the Trinity suite v2.8.5 [44] was run with a k-mer size of 25 and a read coverage normalised to a maximum of 50. In order to minimise the time and computational resources required to analyse and annotate the assembly a number of methods to reduce duplication and filter out sequencing artefacts presenting as lowly expressed transcripts were performed and compared. Presently, the homologous transcripts, with an identity higher than 95%, were clustered by CD-HIT [45] and duplicates removed, in addition the transcripts with an expression lower than 1.5 Transcripts Per Million (TPM) were filtered. The transcriptome assembly quality metrics were measured with the Trinity Perl script “TrinityStats.pl”, the completeness was confirmed using BUSCO against the *Solanaceae* OrthoDB10, containing 3,502 single copy benchmark genes.

In parallel, guided gene prediction with Augustus v3.3.2 [46] was done on the genome assembly. First, RepeatMasker v4.0.9 [47] soft-masked the repeats, using “tomato” as species and a library of repeats, “draft_repeats_master.v5.fasta“, available at ftp://ftp.solgenomics.net/tomato_genome/repeats/. STAR v2.6.0c was used to align the RNA-Seq reads to the masked assembly with intron hints extracted from the resulting BAM file into a GFF file via Augustus’ bam2hints.pl script. Whilst it is possible to extract exon locations to further aid gene prediction, reads may also align to UTRs and thus, due to a lack of trained UTR parameters for *S. sitiens*, this was not performed. Augustus was run using parameters trained for *S. lycopersicum* with repeat masking and intron hints as extrinsic evidence; default hints weighting was used.

Functional annotations of the transcriptome and gene prediction was performed with OmicsBox v1.1.164 [48], using a local BLASTX-fast installation with default parameters against NCBI-nr and custom databases produced from Sol genomics’ *S. lycopersicum* (ITAG3.2) and *S. pennellii* (Spenn) [49] annotated proteins, and The Arabidopsis Information Resources’ (TAIR10) [50]. The blast top-hit results were then associated with Gene Ontology (GO) terms and enzyme annotations. Moreover, protein domains in the transcriptome were identified with an InterProScan [51] search against the following databases, CDD, HAMAP, HMMPanther, HMMPfam, HMMPIR, FPrintScan, ProfileScan and HMMTigr.

For the comparative orthology, protein sequences were extracted from the *S. sitiens* and *S. chilense* gene predictions (unpublished data) GFF output files via Augustus’s Gff2aa.pl script. These protein sequences in addition to sequences from *S. pimpinellifolium, S. lycopersicum, S. pennellii, S. tuberosum* (Sol Genomics) and *A. thaliana* (TAIR10) were entered into OrthoFinder v2.3.3 [52] for the identification of common orthogroups and putative orthologues between each of the species through an all vs all DIAMOND BLAST algorithm. The resulting rooted species tree was plotted using GGtree [53]. Predicted proteins assigned to orthogroups unique to *S. sitiens* were blasted against the NCBI-nr database and functionally annotated facilitating identification of stress-relevant putative proteins.

### Pseudomolecule assemblies based on similar species

The script *chromosome_scaffolder*.*sh* from MaSuRCA was launched to generate two *S. sitiens* pseudomolecule assemblies, based on two different references, *S. lycopersicum v3*.*0* and *S. pennellii*. The pseudomolecules are generated by first splitting the scaffolds into contigs, which are then aligned to the reference sequences with blasr.

For both references, the sequence similarity threshold (-i option) was set to 80, and the PacBio RSII reads were aligned to the reference to check for misassemblies (-s option).

## Results and discussion

### Data quality control

Supplementary Table 5 describes the length distribution of the filtered BioNano molecules and the number of labels/100kbp, which are in the optimal range recommended by BioNano, between 8 and 15 [54].

### The assembly

The statistics obtained at each step of the pipeline are available in Table 1. More details are available in Supplementary Table 1.

Pilon corrected the assembly based on the evidence from the alignments with the Illumina reads, for both libraries >98% of the reads were successfully mapped. Pilon corrected 127,915 single bases, 24,066 insertions amounting to 163,484 bp and 28,397 deletions amounting to 437,669 bp.

GapFiller managed to remove ∼400,000 N’s from the assembly by resizing gaps. There were 18,797,250 N’s remaining after the GapFiller step, corresponding to ∼1.5% of the assembly length.

The blast search of our scaffolds against NCBI-nr matched *Solanum* accessions only. Most of the hits correspond to *S. pennellii*, and 9 scaffolds had no hits, interestingly, these were removed during the deduplication step (see below). The e-value distribution of the hits is available as Supplementary Figure 3A.

**Figure 3:**
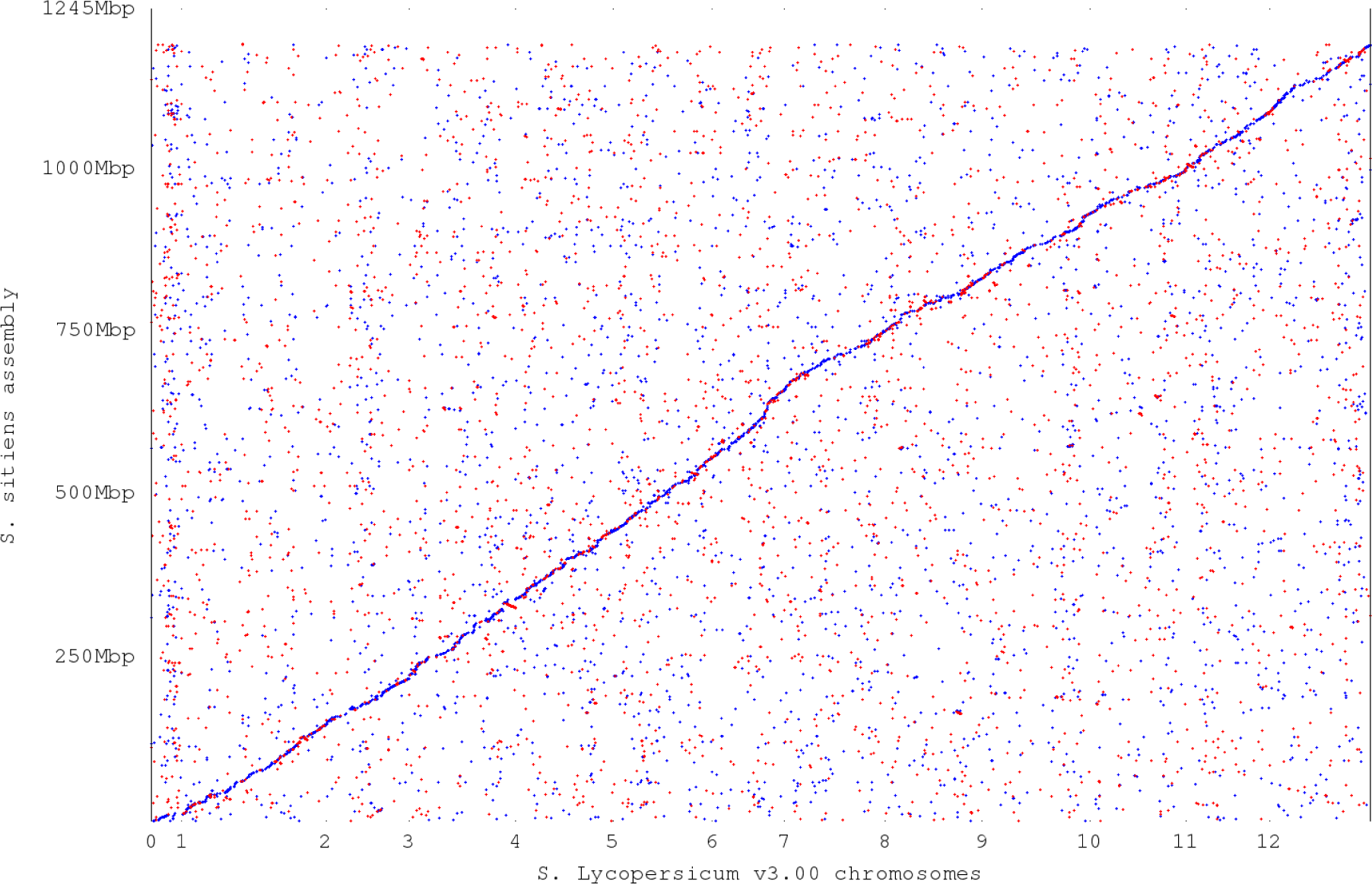
Dotplot of the nucmer alignment between the S sitiens assembly and S. lycopersicum reference genome v3.0, made with Mummer v4. Blue dots represent local forward alignments, red dots represent local reverse alignments.

BBmap dedupe.sh removed 1,202 duplicated scaffolds from the assembly, amounting to ∼29 Mbp. Among these, all the scaffolds without blast hits found previously were removed as duplication during this step. Moreover, most of the scaffolds with a percentage of confirmed bases lower than 95%, as detected by Pilon, were removed (see Supplementary Figure 4) and the average percentage of confirmed bases, increased from 95% to 98%.

**Figure 4:**
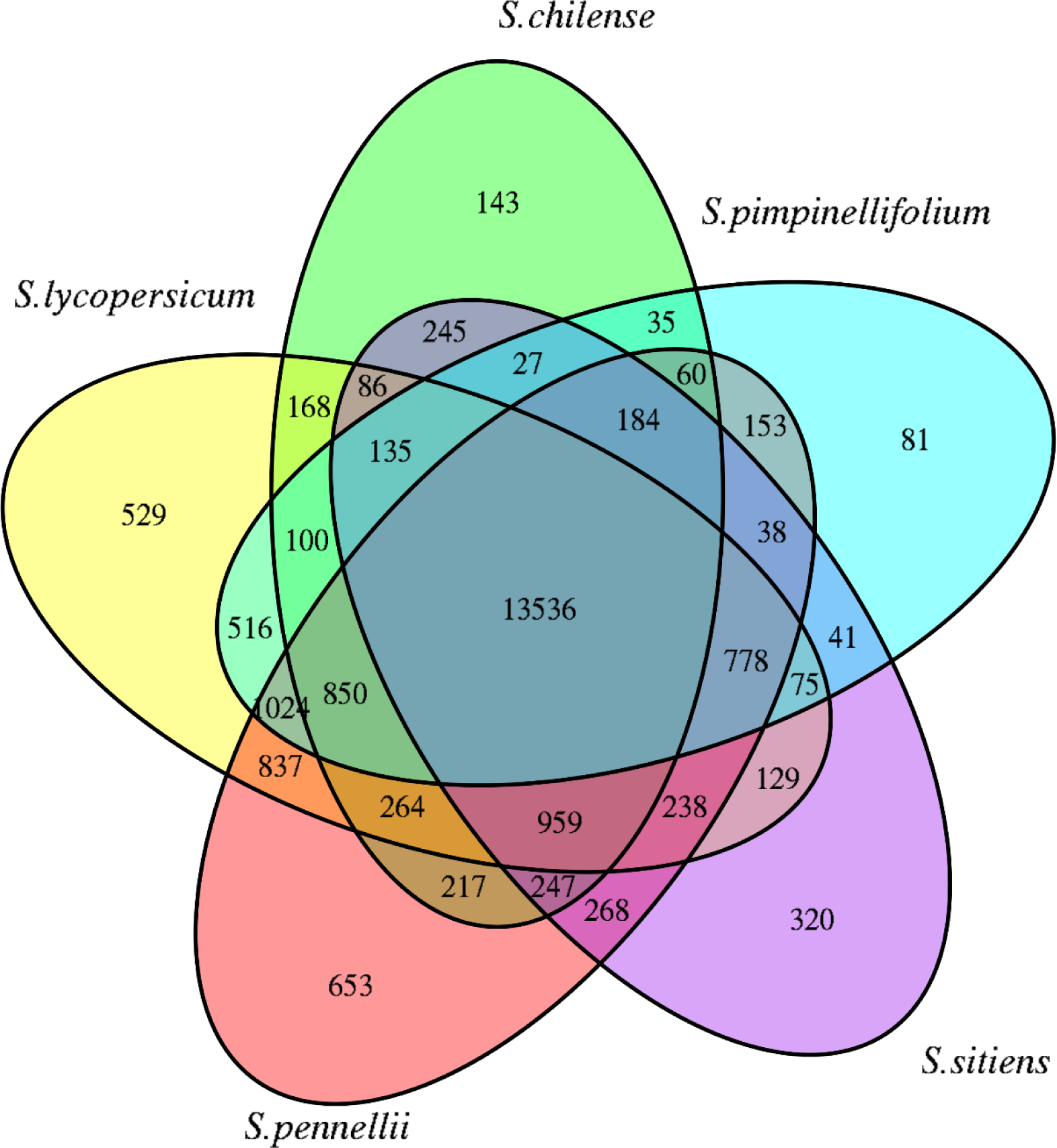
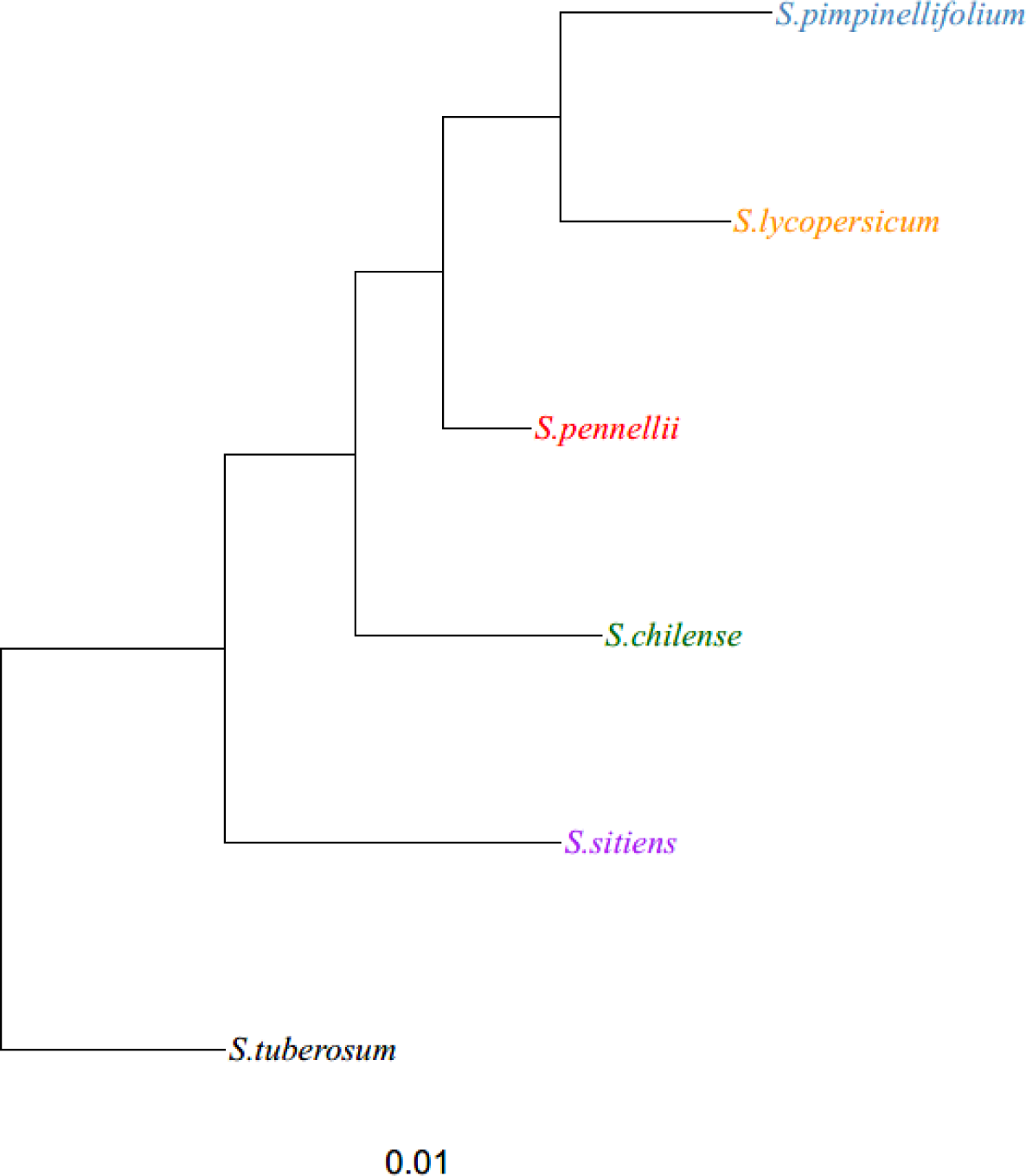
Results from Orthofinder on S. chilense, S. sitiens, S. pennellii, S. lycopersicum, S. pimpinellifolium, S. tuberosum and A. thaliana. **A**. Venn diagram showing unique and shared protein orthogroups assigned to the 5 tomato species. **B**. species tree inferred from the presence of single copy orthologues, produced with GGtree. A. thaliana not represented for clarity’s sake.

### The final assembly quality assessment

The final assembly is composed of 1,481 scaffolds for a total length of 1,245 Mbp, close to the previously estimated genome size of 1,129 Mbp. There are 17,566,158 unknown bases (1.4% of the total length) and 35.34% of the bases are Guanine or Cytosine (GC). The N50 of 1,826,367 bp was found at the 186^th^ scaffold, with the largest sequence being 22,444,909 bp long.

From the 3,052 genes in the Solanaceae dataset, 2,898 (95%) were found in the assembly as complete, this BUSCO result parallels with KAT estimated assembly completeness of 97%, as described below.

KAT estimated a genome size of 841 Mbp, we suspect an underestimation of the size due to KAT ignoring k-mers found outside the heterozygous and homozygous peaks, however, this does not affect the estimation of assembly completeness of 97.28%, which is based on the missing k-mers in the homozygous peak. From the Illumina reads, KAT found three peaks at multiplicity 48, 97 and 192, the first two peaks correspond to the heterozygous and homozygous peaks respectively, and the last peak indicates the presence of regions present in two copies in *S. sitiens* (see Figure 2). The details of KAT output are available in Supplementary Table 8.

The overall mapping rates of the 4 RNA-Seq libraries against the assembly were between 91.91% and 94.86%. Similar to the approach taken by *Bolger et al*. [13] the unmapped reads were re-mapped to *S. lycopersicum* v3.0 to determine if they were due to assembly incompleteness, only 1.32% to 1.76% of the reads mapped to *S. lycopersicum* and not our assembly, indicated therefore no loss of data due to assembly incompleteness.

### Comparison with closely related assemblies

The genome assembly overall structure and completeness was assessed further by performing an overall pairwise alignment against *S. lycopersicum* and *S. pennelliii*. Figure 3 shows the alignment of our *S. sitiens* scaffolds against all twelve chromosomes of *S. lycopersicum*. It can be noted that the developed *S. sitiens* assembly is showing an overall continuity across all chromosomes. Similar pairwise alignment against *S. pennellii* is also available as Supplementary Figure 5 and similar coverage was also obtained.

Statistics of our MaSuRCA assembly were compared against the contig assemblies of *S. pennellii, S. pimpinellifolium* and *S. lycospersicum* v3.0, see Table 2. There were obtained with Quast using the “--scaffold” option to break the scaffolds by stretch of Ns longer than 10. This allowed fair comparison between the assemblies at the scaffold and chromosome level.

A BUSCO analysis, on the original fasta files, allowed comparison between our assembly and the assemblies of *S. lycopersicum* v3.0, *S. pennelllii, S. pimpinellifolium*. See Supplementary Table 10 for more details on the Busco results.

### De novo transcriptome assembly and functional annotation

Quality assessment for raw reads was performed using Rcorrector. 8.4% of the RNA-seq reads were identified as ‘unfixable’; and were marked as such should the number of detected errors across the length of the read exceed a pre-determined threshold. These unfixable reads may either be derived from a lowly expressed transcript of which a more greatly expressed homolog is present, or may simply contain too many errors to be corrected. To avoid the inclusion of damaging k-mers, all unfixable reads were removed. The error correction of the reads with low quality did not remove any additional reads, but trimmed 1.62% of the base pairs by cutting off the adapters. Ultimately, the cleaned data retained in average 32 million paired-reads per sample.

The raw transcriptome from Trinity generated 461,083 contigs for a total size of 387 Mbp and an N50 of 1,710 bp. This is obviously much higher than the expected number of transcripts and is usually the case with unfiltered output from Trinity. Moreover, high levels of duplication are common and to be expected in transcriptome assembly due to the transcription of multiple isoforms of the same RNA-product. Therefore, duplicates and lowly expressed transcripts were considered as artefacts and were subsequently removed. CD-HIT clustered similar sequences and removed 118,638 duplicates. The transcriptome was filtered by a threshold of 1.5 TPM, which was the optimum compromise between duplication level, transcript count and completeness. The final transcriptome contains 158,918 transcripts, with an estimated BUSCO completeness of 90.0%. More details about the *de novo* transcriptome assembly statistics are available in Supplementary table 11.

The functional annotation was done by aligning the transcriptome against NCBI-nr and custom blast databases produced from the assemblies of closely related species. During the IPS search, 77,834 transcripts had blast hits, including 35,645 transcripts associated with GO terms. InterproScan repeats distribution results showed Leucine-rich repeat (LRR) proteins repeating 456 times. This domain is directly involved in recognising the presence of pathogen-associated molecular patterns (PAMPs). The pathogen products of avirulence (AVR) genes are recognised by the nucleotide-binding site of LRR proteins. A gene coding the protein with Ankyrin repeats domain found 169 times in InterproScan results shows increased resistance to disease and spontaneous cell-death. Spontaneous cell death is a critical disease response mechanism where a plant kills the diseased cells instantly to block the disease from spreading. An in-depth look at how and when these genes are over-expressing in *S. sitiens* can provide more understanding of its response to pathogens and diseases.

### Gene prediction from the genome assembly

Excluding gaps from the assembly, RepeatMasker masked 70% of the genome as repetitive elements, which is in the range of other *Solanum* accessions, eg. 59.5% for *S. pimpinellifolium* [17], 64% for *S. lycopersicum* v4 [55], and 82% for *S. pennellii* [13].

Augustus predicted 31,164 genes, corresponding to 33,386 transcripts, using hints about the intron positions obtained from the alignment of the RNA-seq against our genome assembly. This number is very close to the number of coding genes present in *S. lycopersicum* (34,658) and *S. pennellii (*32,273).

### Comparative orthology reveals

Orthofinder revealed 23,225 orthogroups in total between *S, sitiens, S. chilense, S. pennnellii, S. pimpinellifolium, S. lycopersicum, S. tuberosum* and *A. thaliana*, defined as the set of genes descended from a singular gene of the last common ancestor of each species. A rooted phylogenetic tree was inferred through the presence of the most closely related genes within single and multi-copy orthogroups (see Figure 4B). As expected *S. pimpinellifolium* and *S. lycopersicum*, which present only 0.6% nucleotide divergence and recent introgressions [21], are shown to be the most closely related of the tomato species whereas the potato, *S. tuberosum*, is the least closely related species to the domestic tomato. The correct placement of these three species lends credence to the accuracy of the placement of *S. sitiens* and *S. chilense* within the phylogram and the output of Orthofinder. Of the five tomato species considered, 13,536 orthogroups were common to all species (see Figure 4A). A significant number of orthogroups were found to be unique to each species, evidence of the genetic diversity between tomatoes. However, it should be noted that individual tomatoes will harbour different genes as revealed through the investigation of the tomato pan-genome [21] and thus the extent of unique orthogroup assignment is to a certain extent dependent on the individuals from which the protein sequences are derived.

### Annotation of the transcripts unique to *S. sitiens*

Some of the 561 transcripts belonging to the 320 orthogroups unique to *S. sitiens* (see Figure 4A) have annotations related to salt and drought stress tolerance. Notably, *Glycolate Oxidase* (*GLO1* and *GLO4*), which are involved in drought stress response in *Vigna* [56] and pea [57]. One of the transcript mapped with the GO term “response to water deprivation” (GO:0009414) and blasted against an aquaporin *PIP1-2* which plays a role in drought tolerance [58]. Another transcript mapped to the GO term “response to water” (GO:0009415) and blasted against “abscisic acid and environmental stress-inducible protein *TAS14*” which was found to improve both salt and drought resistance [59]. Other interesting transcripts blasted against *Oxophytodienoate Reductase 1* and *FQR1-like NAD(P)H dehydrogenase* which were shown to confer salt tolerance in wheat [60] and Arabidopsis [61] respectively. Very interestingly, genes with high homology to *FQR1-like NAD(P)H dehydrogenase* were also identified in *S. pimpinellifolium*, another drought and salt tolerant tomato wild relative [17]. Further study of these unique orthologs might give hindsight into the adaptation of *S. sitiens* in water limited environments.

### Identification of inversions against *S. lycopersicum*

Three potentially interesting inversions against *S. lycopersicum* were identified using the alignments produced with Mummer (see Supplementary Figure 6). One is on scaffold95 against chromosome 11 of *S. lycopersicum* and is present both in our assembly and *S. pennellii*. This inversion has also been previously reported in *S. habrochaites* against *S. lycopersicum* [62]. The other two, on scaffold11 and scaffold8 are unique to *S. sitiens* against both *S. lycopersicum* and *S. pennellii*. All three inversion were located within contiguous sequences on our scaffolds. The locations of these inversions were intersected with the gene prediction from Augustus. The genes found in the inversions were blasted against ITAG4.0 and TAIR10 and the top hits were searched in the literature for links to drought or salt resistance (see Table 3). The Arabidopsis orthologs obtained from the literature search were then queried against the ePlant web repository [63] to look for expression changes during drought and salt stresses (“Abiotic Stress eFP” and “Abiotic Stress II eFP” views under “Tissue & Experiment eFP viewers”) [64] [65].

**Table 3:** List of genes related to drought and salt tolerance in the inversions against S. lycopersicum. The genes were obtained from the gene prediction done on the assembly with Augustus. The Solyc ID were obtained by blasting the predicted gene against the ITAG 4.0 annotation (ITAG4.0_gene_models.gff). The distance to the nearest breakpoint is an estimate based on the dotplots, the genes located outside an inversion have a negative value. **A**. Inversion on Scaffold95 **B**. Inversion on Scaffold8 **C**. Inversion on Scaffold11

| Scaffold95:800000 - 1120000 |  |  |  |
| --- | --- | --- | --- |
| Blast hit (ITAG4.0) | Distance to nearest breakpoint (kbp) | Gene name | Literature |
| Solyc11g069960 | -31 | receptor-like kinase RLK-1 | *RLK1* is involved in response to salinity and drought stresses by activating or participating in ABA signaling [66] [67]. |
| Solyc11g069940 | -5 | Glutaredoxin | In tomato, *SIGRX1* silencing increased the plant sensitivity to both drought and salt stresses [68]. In cassava cultivars and maize, *CC type glutaredoxins* have been shown to be associated with drought response [69] [70]. |
| Solyc11g069690 | 32 | protein disulfide-isomerase 5-1 | In *A. thaliana*, *PDI5* is involved in drought signaling [71]. Moreover, members of this protein family (*PDI*) are improving protein folding and transport during stress [72]. |
| Solyc11g069700 | 42 | elongation factor 1-alpha | In soybean, *EF1α* genes were upregulated by ABA, salt and drought stresses [73]. |
| Solyc11g069735 | 80 | High-affinity nitrate transporter 2.2 | Higher expression of *NRT2.1* and *NRT2.2* was revealed to enhance drought resistance in *A. thaliana* [74]. |
| Solyc11g069765 | 111 | SNF1-related protein kinase regulatory subunit beta-3 | In plants, some members of the SNF1/AMP-activated kinases are activated by dehydration or ABA [75]. |
| Solyc11g069770 | 130 | agamous-like MADS-box protein AGL62 | Considered as a top candidate gene for a "Water Use Efficiency" in dry condition QTL in *A. thaliana* [76]. |
| Solyc11g069800 | 123 | Allene oxide synthase, chloroplastic | AOS is an important enzyme in the jasmonic acid pathway, which plays a role in the plant response to salt stress. AOS accumulated in salt tolerant tomato plants after 6 hours of NaCl and jasmonic acid treatment [77]. In sweet potato and Arabidopsis, AOS was upregulated under salt stress [78] [79]. |
| Solyc11g069820 | 45 | ABC transporter-like | Members of this family annotated with GO terms “response to salt stress” (GO:0009651) and “response to abscisic acid” (GO:0009737).<br>Overexpression of *AtABCG36* led to improved drought and salt tolerance in *Arabidopsis* [80]. |
| Solyc11g069910 | 21 | DNA-directed RNA polymerase II subunit RPB11 | In *Arabidopsis*, RNA polymerase II quickly acted on drought-responsive genes when subjected to drought stress and rapidly disappeared during rehydration [81]. |

B.
| Scaffold8:2250000-2430000 |  |  |  |
| --- | --- | --- | --- |
| Blast hit (ITAG4.0) | Distance to nearest breakpoint (kbp) | Gene name | Literature |
| Solyc09g091170 | -8 | NADH dehydrogenase | In *Nicotiana tabacum*, two NADH dehydrogenases are over expressed during drought stress [82]. *AtNDB2*, an *Arabidopsis* NADH dehydrogenase was identified as being important for tolerance to environmental stress. Plants lacking *atNCB2* were less tolerant to drought and elevated light treatments [83]. |
| Solyc09g091030 | 19 | Beta-amylase | *BAM1* is annotated with the GO Term “response to water deprivation” (GO:0009414).<br>*BAM1* is involved in the starch/proline interplay, which has been identified as a candidate trait to improve drought tolerance in crops [84]. Drought tolerance was improved in an *A. thaliana* *BAM1* mutant, because the resulting decrease in starch breakdown led to reduced stomatal opening [85]. |
| Solyc09g091060 | 44 | golgin subfamily A member 6-like protein 2 | This gene has been reported as up-regulated in the leaves of UCB-1 pistachio rootstock under drought stress, with a probable role in transportation and glycosylation of lipids and proteins under water deficit [86]. |
| Solyc09g091090 | 66 | probable indole-3-pyruvate monooxygenase YUCCA7 | *YUCCA7* is annotated with the GO Term “response to water deprivation” (GO:0009414).<br>From the TAIR10 description (AT2G33230.1): “Encodes a flavin monooxygenase gene which belongs to the tryptophan-dependent auxin<br>biosynthetic pathway and enhances drought resistance.”<br>In *Arabidopsis*, activation of *YUCCA7* led to increased drought tolerance [87]. In rice, this gene is involved in auxin production, which increase root development and makes the plant more adapted to drought conditions [88]. |
| Solyc09g091120 | 47 | BLISTER | *BLISTER* was shown to promote resistance to cold and drought stresses, by repressing ABA-reponsive PcG target genes in *A. thaliana* [89]. |
| Solyc09g090980 | -3 | pathogenesis-related protein STH-2-like (alternative name: PR10) | *SmPR10* expression in transgenic *Arabidopsis* was up-regulated, specific to roots and improved the plants tolerance to salt stress [90]. |
| Solyc09g090920 | -29 | Calcium-dependent lipid-binding (CaLB domain) family protein | In *A. thaliana*, loss of function of *AtCLB* increased the plant's tolerance to salt and drought stresses [91]. |

C.
| Scaffold11:5030000-5160000 |  |  |  |
| --- | --- | --- | --- |
| Blast hit (ITAG4.0) | Distance to nearest breakpoint (kbp) | Gene name | Literature |
| Solyc10g079750 | -33 | BTB/POZ ankyrin repeat protein | *BOP1* and *BOP2* are necessary for cauline leaf abscission in response to drought in *Arabidopsis* [92]. |
| Solyc10g079600 | 8 | Two-component response regulator ARR9 | In *Arabidopsis*, *ARR9* was suggested to have a complex role, both positive and negative, in osmotic stress regulation [93]. |
| Solyc10g079580 | -5 | RNA-binding (RRM/RBD/RNP motifs) family protein | RNA-binding mechanisms might have roles in drought and salt stress response [94] [95]. |

As mentioned in the introduction, *S. sitiens* possess a unique fruit maturation process. Instead of ripening, the fruit are desiccating allowing the seeds to disperse in the desert. Interestingly, one of the genes in the inversion on scaffold11, *ARR9*, is involved in transcription of ABA biosynthetic genes, which in turn is affecting seed desiccation tolerance [93].

The most promising inversion is the one between scaffold8 and chromosome 9 of *S. lycopersicum*. It contains two genes annotated with the GO Term: “response to water deprivation”, *YUCCA7* and *BAM1*, making it a locus of interest for drought tolerance.

Some Arabidopsis orthologs of the genes identified in Table 3 change their expression during drought and salt stresses, as shown on ePlant. Notably, *allene oxide synthase* and *BAM1* are up regulated in both drought and salt stresses. *YUCCA7* is following the same trend except in roots, were it is down regulated during salt stress. *ABC transporter* is up regulated during salt stress. On the contrary, protein *disulfide-isomerase 5-1* and *ARR9* are down regulated during salt stress. *High-affinity nitrate transporter 2*.*2* seems to undergo a lot of change in expression during both drought and salt stresses. The ePlant figures are available as Supplementary Figure 8.

We hypothesise that these inversions might affect the expression of the genes described in Table 3 and renders the plant more susceptible to drought and salt stresses. Further research will be needed to confirm and understand the interplay between these inversions and S. *sitiens* adaptation to its environment.

### Pseudomolecule assemblies based on similar species

The *chromosome_scaffolder,sh* script from MaSuRCA, produced two *S. sitiens* chromosome scale assemblies, one based on *S. lycopersicum*, the other on *S. pennellii*. The assemblies’ statistics and quality were assessed with the same tools and parameters as the scaffold assembly and the results are available in *Table 4*. 81% and 83% of the total length was covered across 12 chromosome sequences for the assemblies based on *S. lycopersicum* and *S. pennellii* respectively. Moreover, the pseudomolecules assemblies were aligned with mummer against their respective references, with the same parameters as described in the methods section “Assessment of the assembly quality“. The dotplots generated from the alignment are available as Supplementary Figures 7A and 7B.

**Table 4:** Pseudomolecule assemblies statistics.

| Assembly name | Scaffold assembly | Pseudomolecule assembly based on *S. lycopersicum* 3.0 | Pseudomolecule assembly based on *S. pennellii* |
| --- | --- | --- | --- |
| Length (Mbp) | 1,245 | 1,703 | 1,822 |
| N50 (bp) | 1,826,367 | 113,281,409 | 123,136,548 |
| Largest scaffold (bp) | 22,444,909 | 174,954,540 | 186,599,541 |
| # scaffolds | 1,481 | 3,042 | 2,877 |
| N / 100 kbp | 1,410 | 27,964 | 32,724 |
| KAT estimated completeness | 97.28% | 97.25% | 97.22% |
| BUSCO completeness | 94.9% | 94.9% | 94.9% |

While these pseudomolecules assemblies are not as accurate as if they were constructed purely from *S. sitiens* data, due to some potential misassemblies stemming from the differences between the genomes of *S. sitiens* and the two relative species used as references. Nevertheless, we decided to release these assemblies as they can be beneficial for genotyping and visualisation purposes via genome browsers.

## Conclusion

Here, we present high-quality *de novo* genome and transcriptome assemblies of *S. sitiens*, a tomato wild relative. This scaffold assembly is more than 95% complete, as measured by BUSCO and KAT. Comparison at the contig level shows better contiguity than assemblies of similar *Solanaceae* species, which will facilitate consequent analyses, notably gene prediction, detection of structural variations and pseudomolecule assembly. Analysis of *S. sitiens* unique orthologs and three inversions against *S. lycopersicum* highlighted genes that could be involved in drought and salt tolerance, this will lead the way for future discoveries on the importance of these genes. Moreover, we are hoping that the availability of this reference will help the breeding efforts to integrate drought resistance in tomato crops.

## Data availability

*S. sitiens* (LA1974) raw sequencing, transcriptome and genome assembly have been deposited at the NCBI’s Sequence Read Archive, under the BioProject number “PRJNA633104”.

### Glossary

ABA: Abscisic Acid
bp: basepair
bwa: Burrow-Wheeler Aligner
BUSCO: Benchmarking with Universal Single-Copy Orthologs
FR: Forward Reverse
Gbp: Gigabasepair
GC: Guanine-Cytosine
GO: Gene Ontology
IPS: InterProScan
KAT: K-mer Analysis Toolkit
Kbp: Kilobasepair
Mbp: Megabasepair
PacBio: Pacific Biosciences
PE: Paired-End
QTL: Quantitative Trait Locus
SMRT: Single Molecule, Real-Time
TAIR: The Arabidopsis Information Resource
TPM: Transcripts Per Million
WGS: Whole Genome Sequencing

