## Supplemental Figures for "*De novo* genome assembly and transcriptome analysis for the drought and salt resistant *Solanum sitiens*"

**A.**
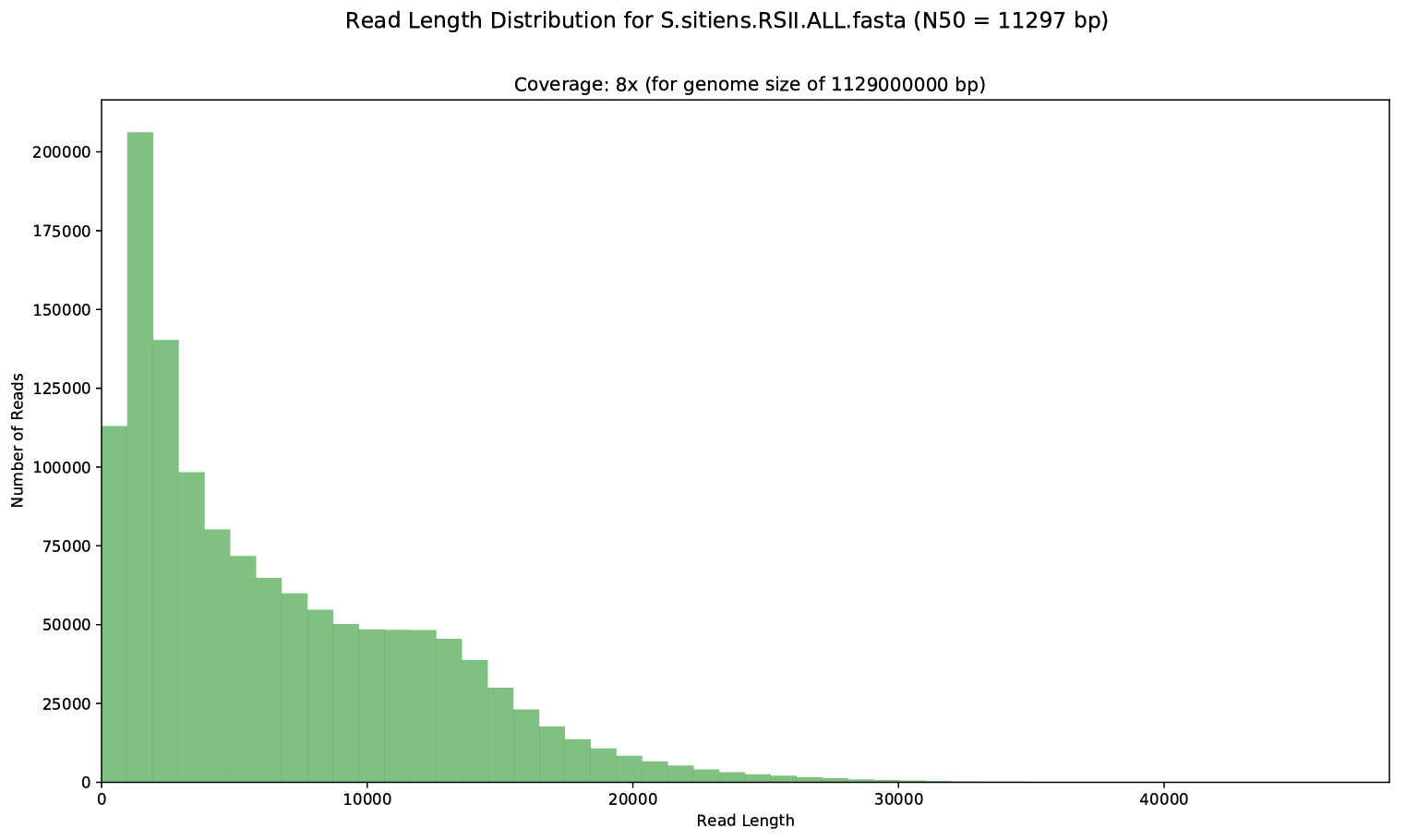

**B.**
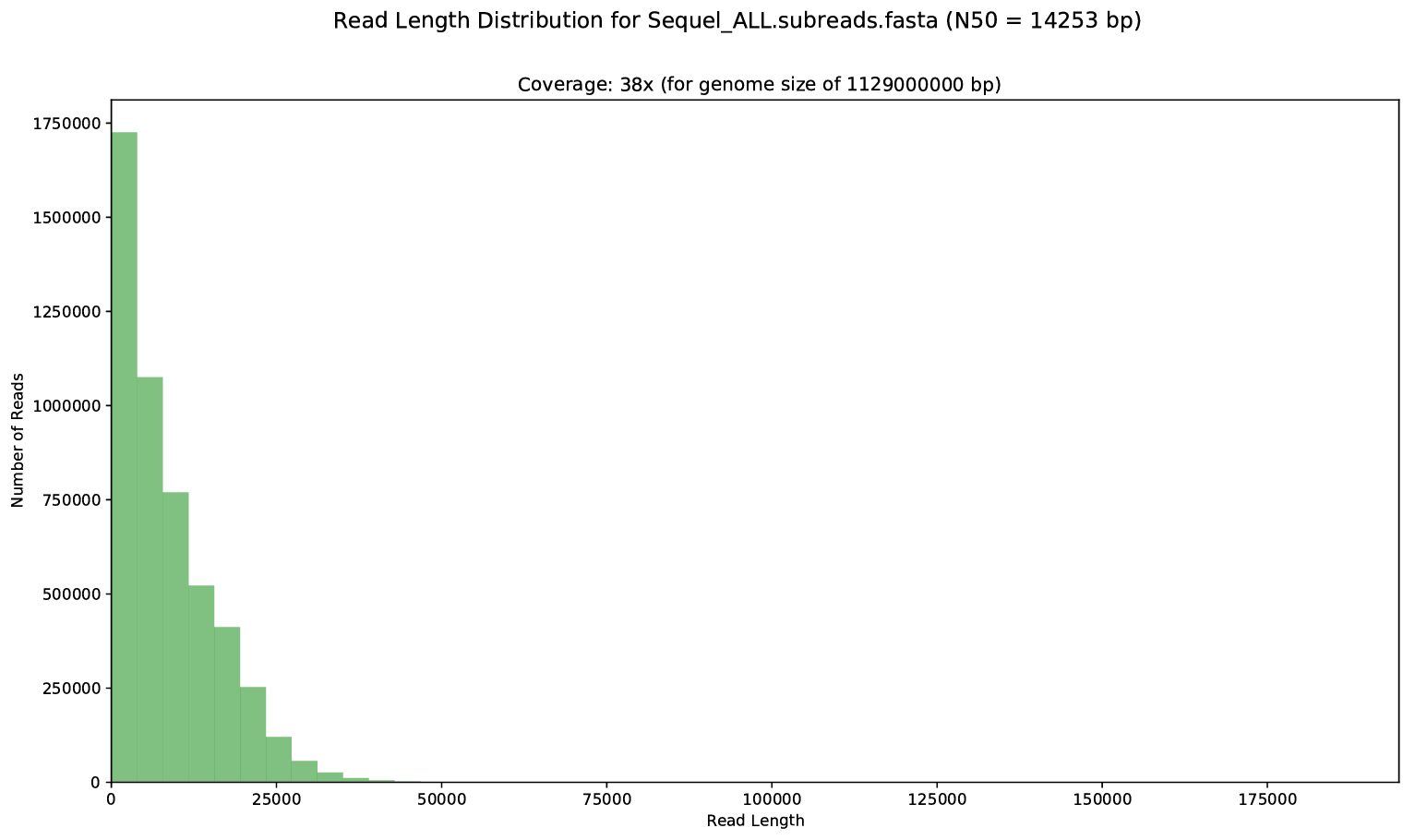

Supplementary Figure 1: Length distribution of the Pacbio reads, filtered by length lower than 1 kbp **A**. the RSII platform **B**. the Sequel platform

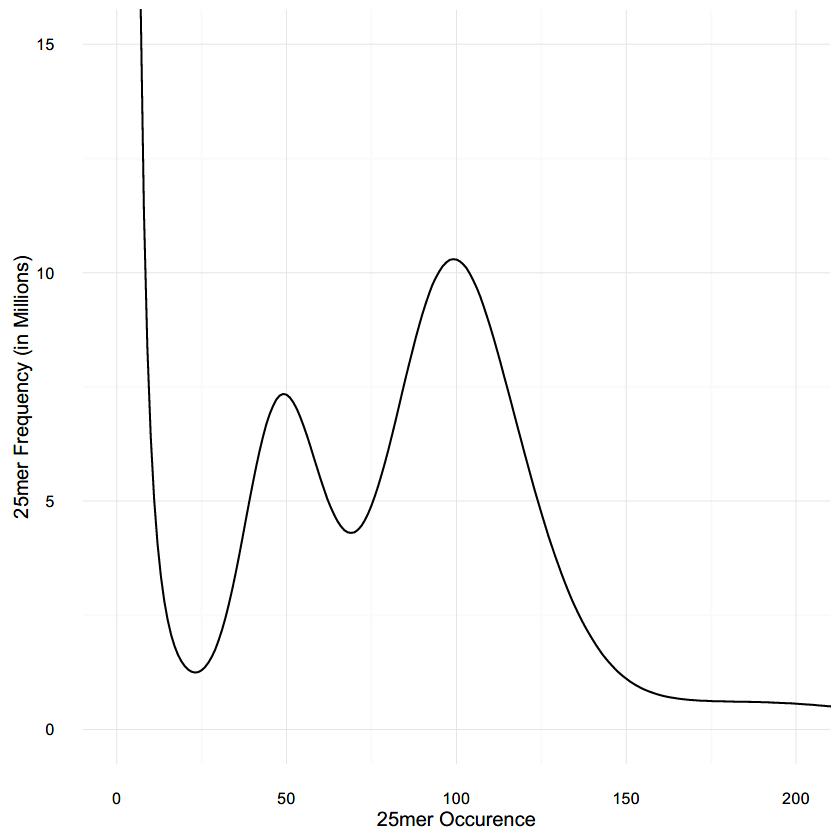

Supplementary Figure 2: Histogram of the 25-mer distribution in the Illumina reads. Two peaks are visible, one at 49 the other at 99, corresponding to the heterozygous and homozygous peak respectively. The low frequency 25-mers, below 24, are considered artefacts.

**A.**

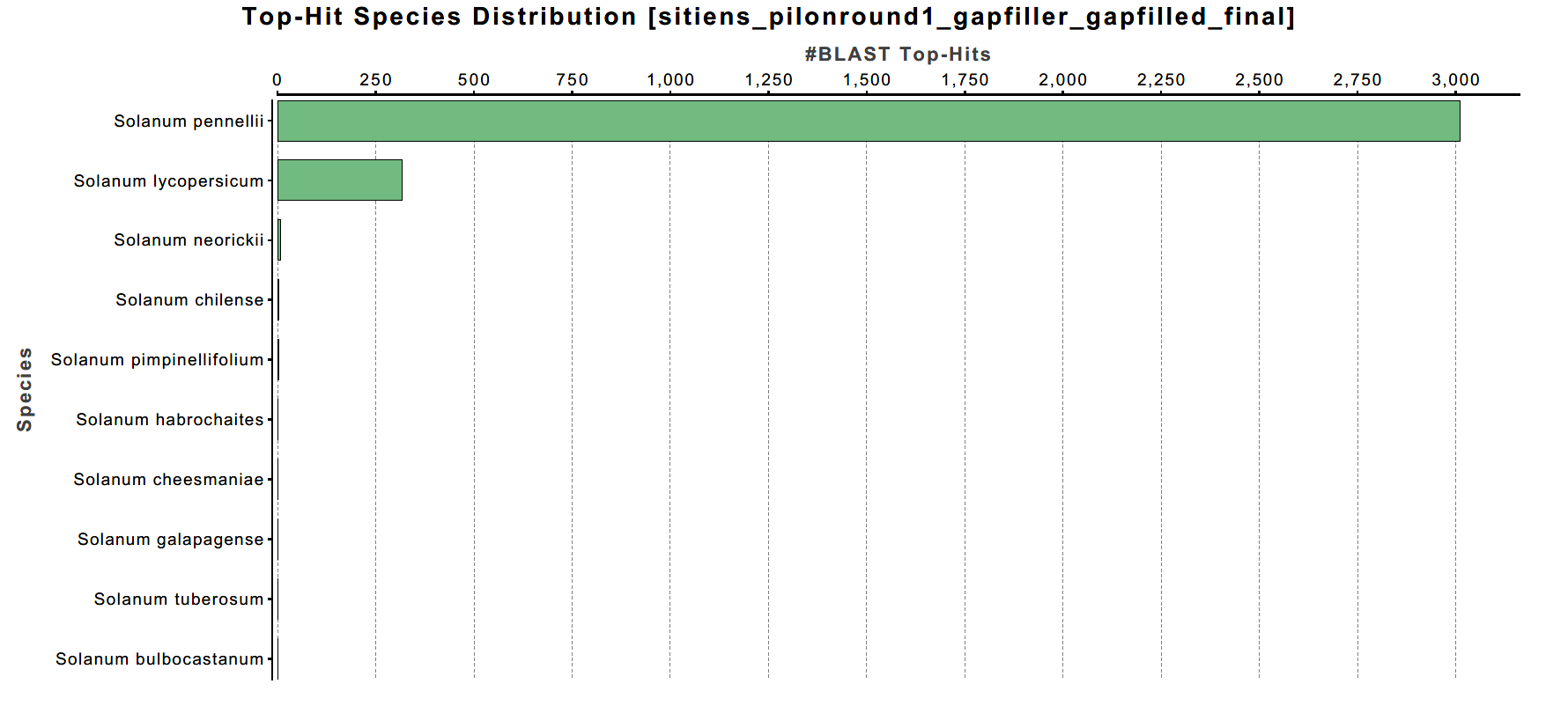

**B.**

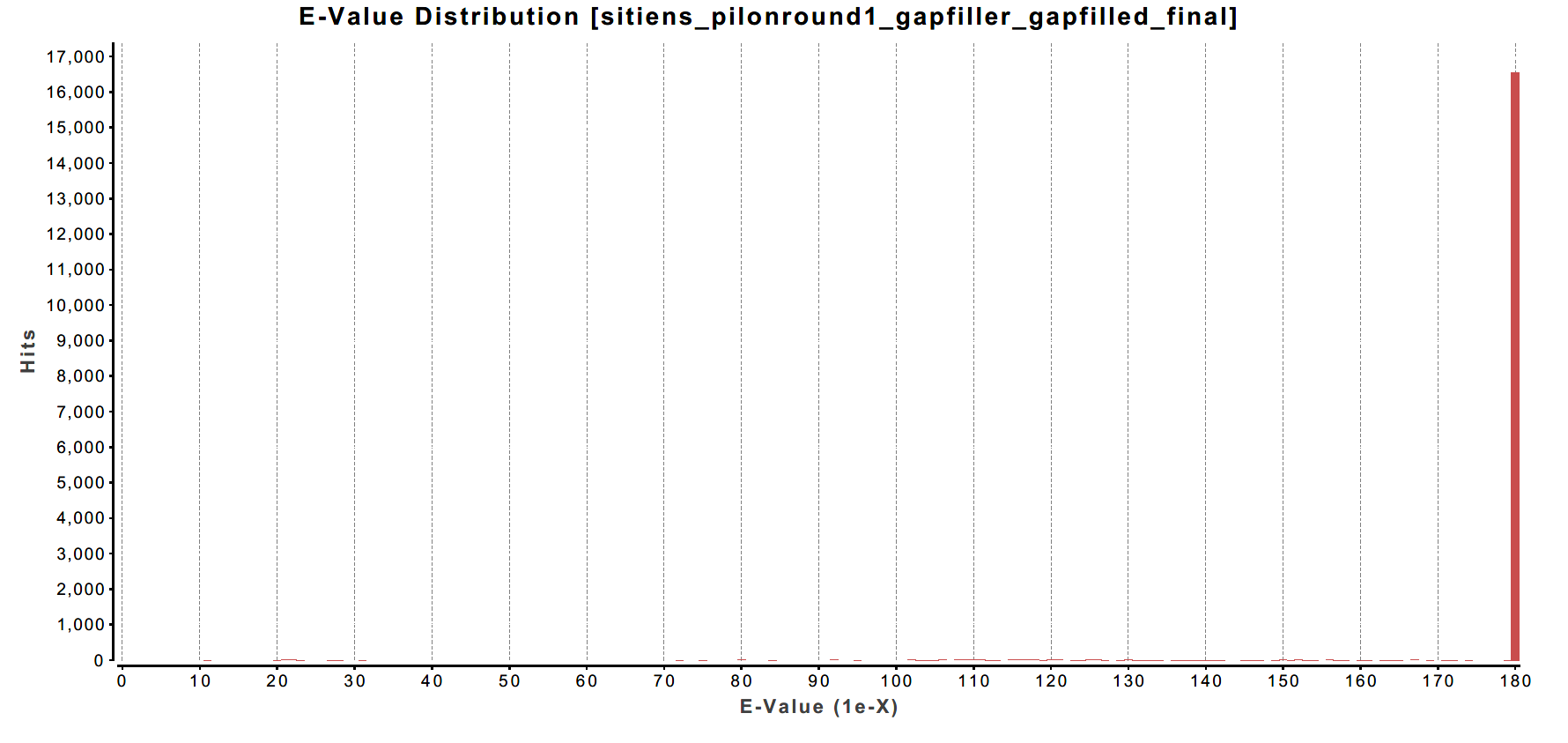

Supplementary Figure 3: Results from the blast search of the scaffolds against NR. **A.** Distribution of the species for the blast top hit, consisting only of Solanum species as expected. **B.** Distribution of the e-value for all the hits, most of the hits have an e-value < 1e-180.

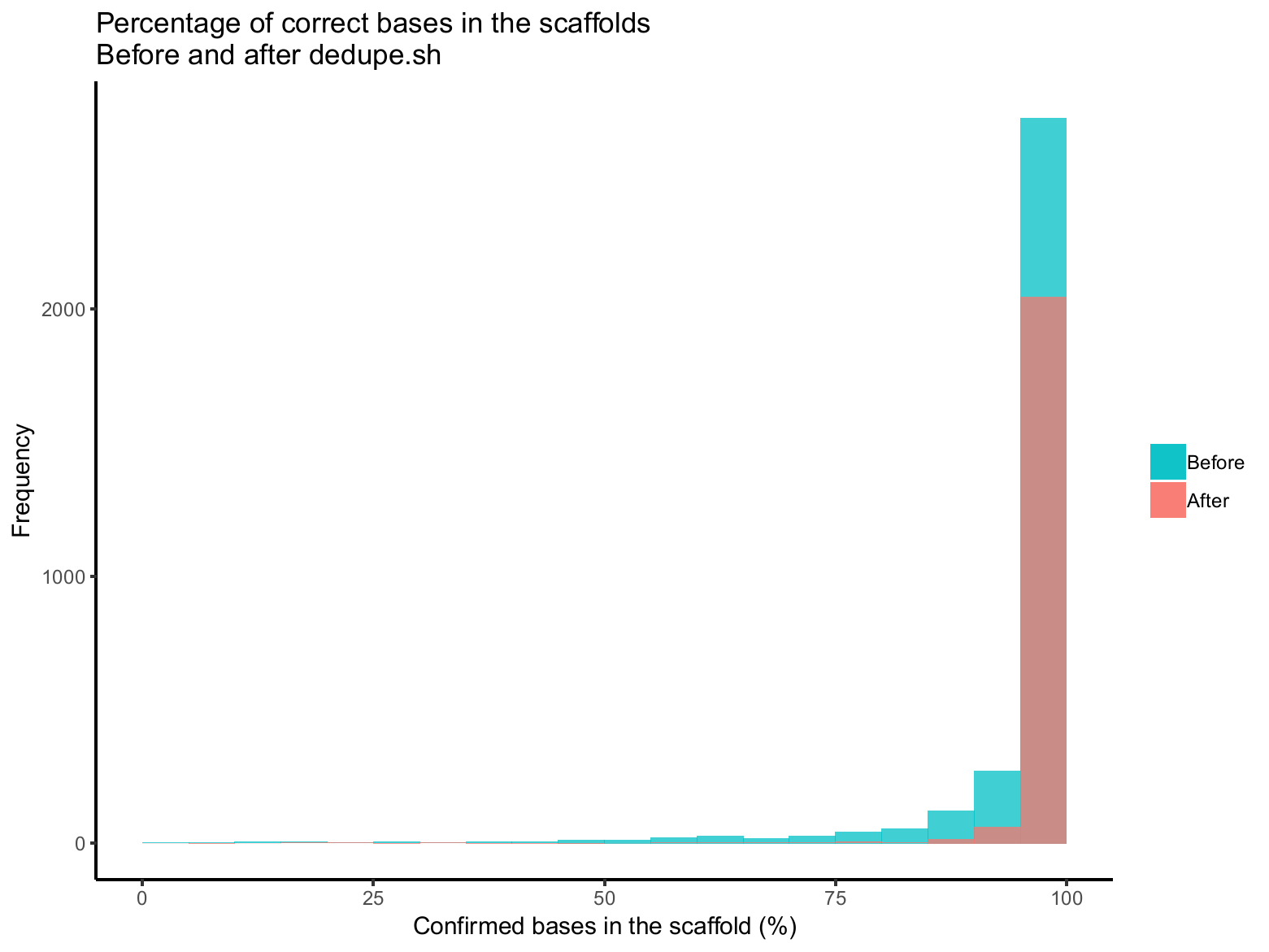

Supplementary Figure 4: Pilon estimation of the correct bases in the assembly, based on the Illumina reads, before and after dedupe. The x-axis represents the percentage of correct bases, the y-axis represents the number of scaffolds. Most of the scaffolds with a percentage of correct bases lower than 95% were removed by dedupe. These were probably low quality duplications due to S. sitiens heterozygous nature. After dedupe, the number of sequences with 100% confirmed bases is also lower, this is because the small contigs corresponding to alternative haplotypes have been removed, as expected in an assembly.

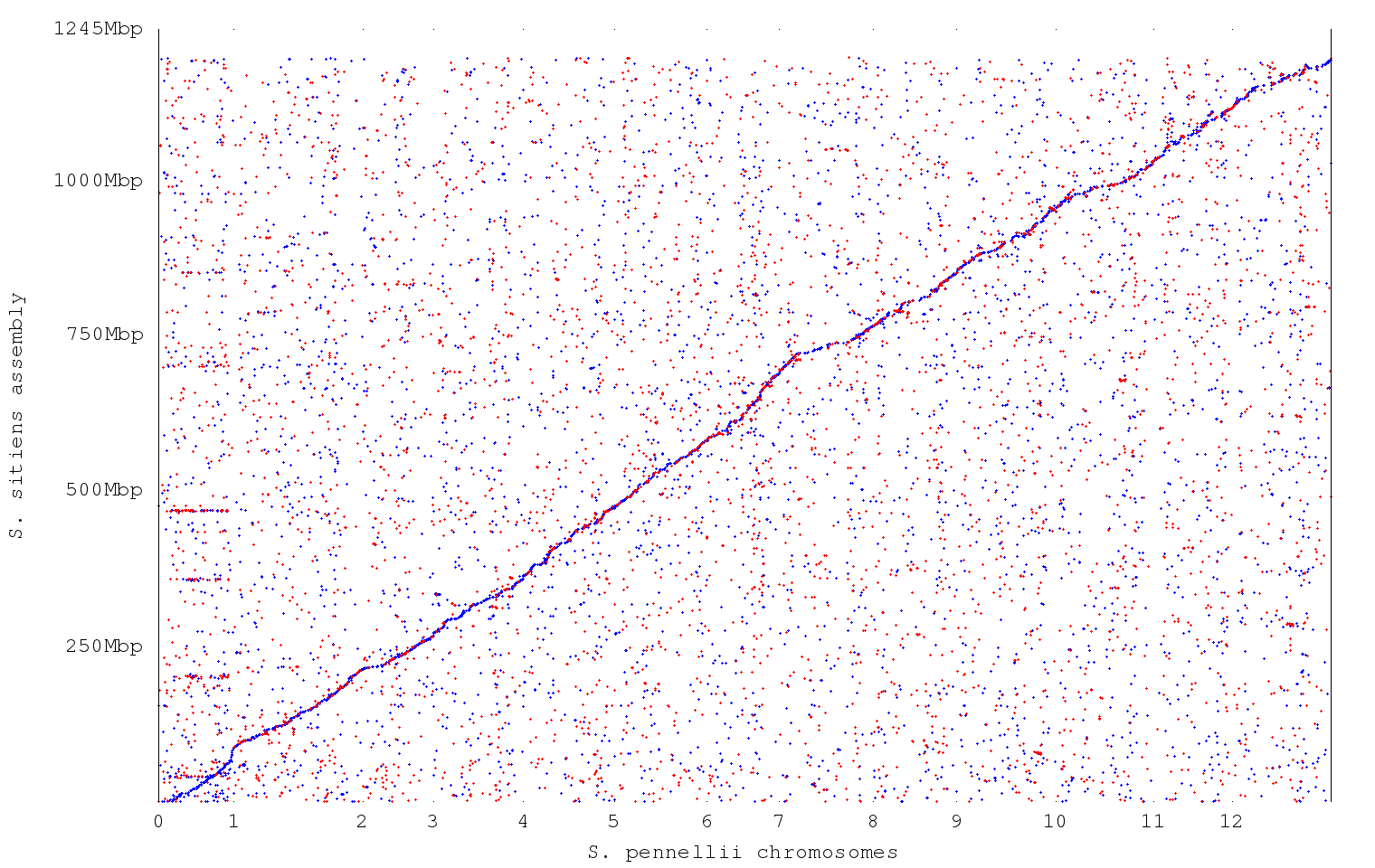

Supplementary Figure 5: Dotplot of the similarity between the S sitiens assembly and the reference genome of S. pennellii, made with Mummer v4.00. Blue dots represent local forward alignments, red dots represent local reverse alignments.

**1A.**

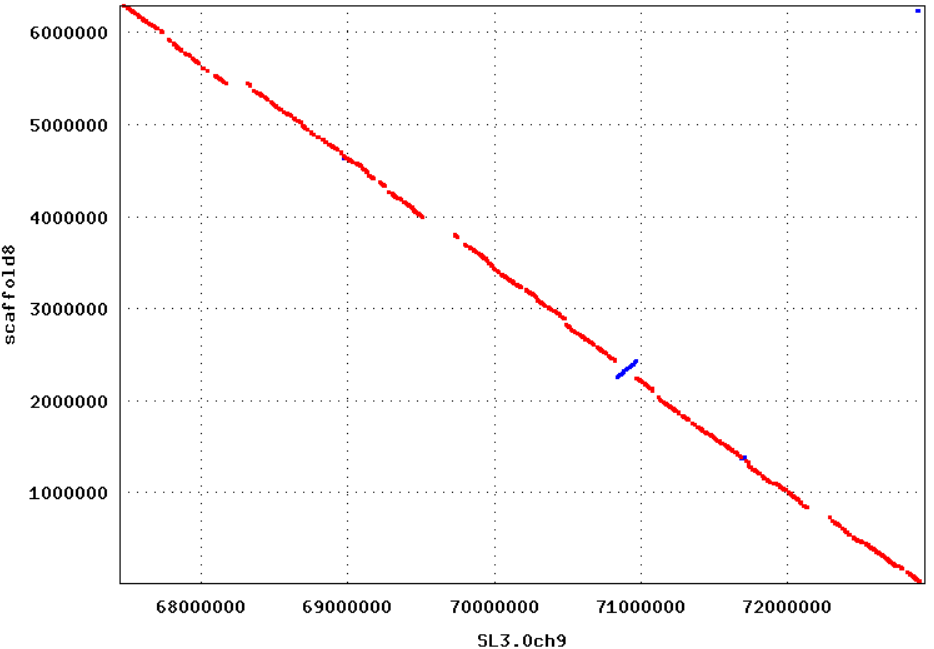

**1B.**

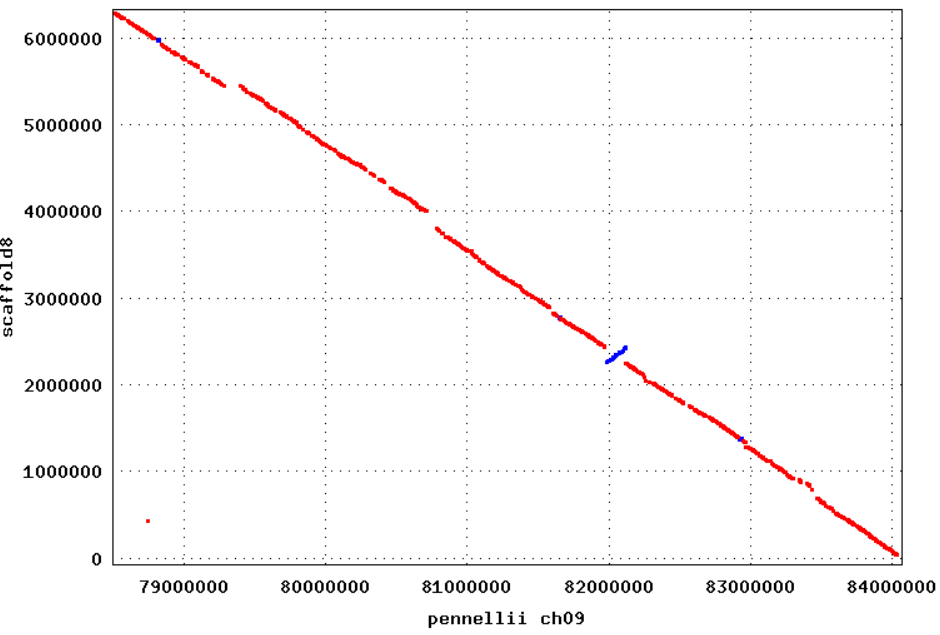

**2A**
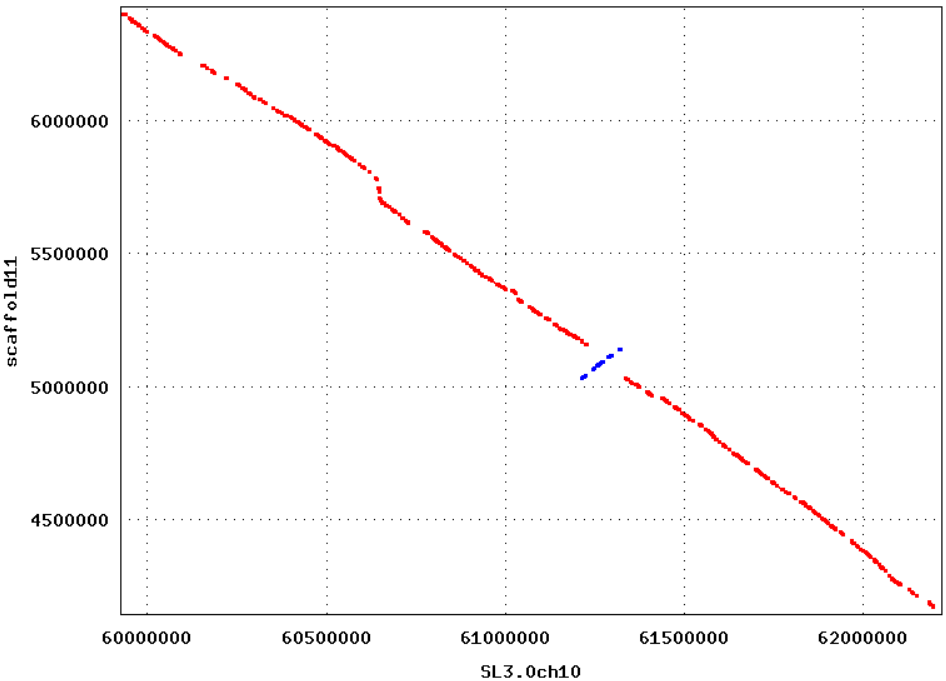

**2B.**
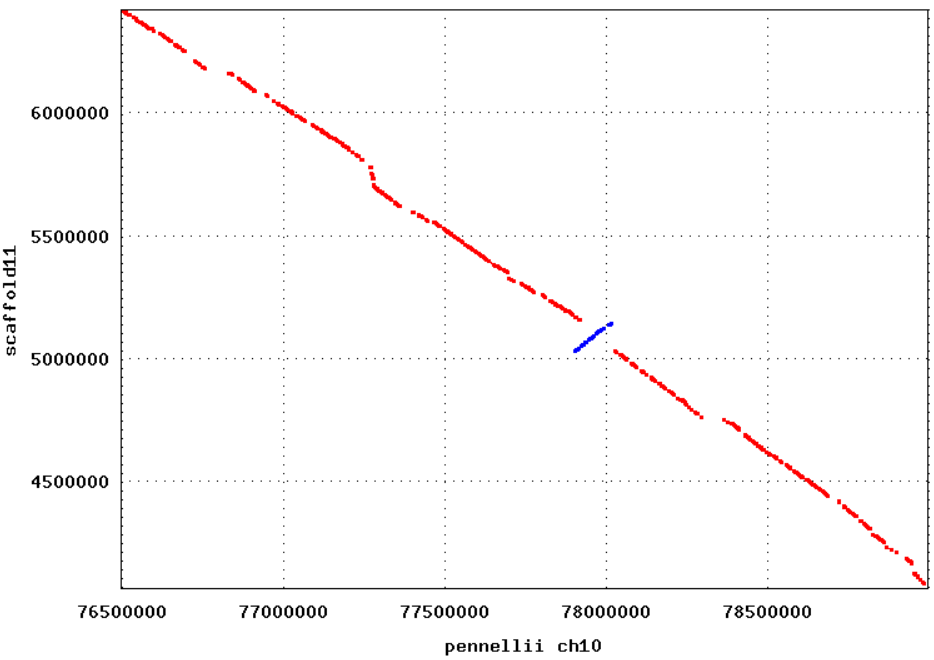

**3A.**

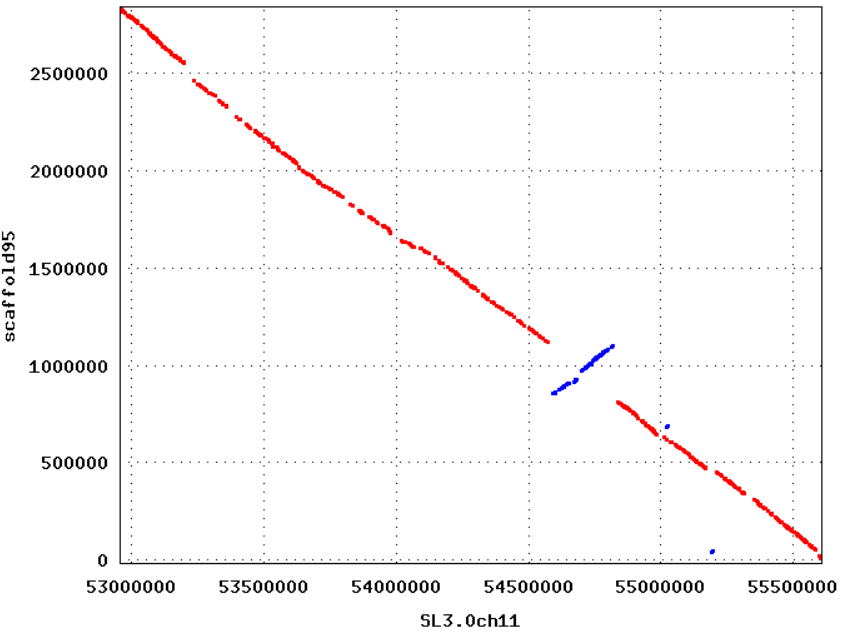

**3B.**
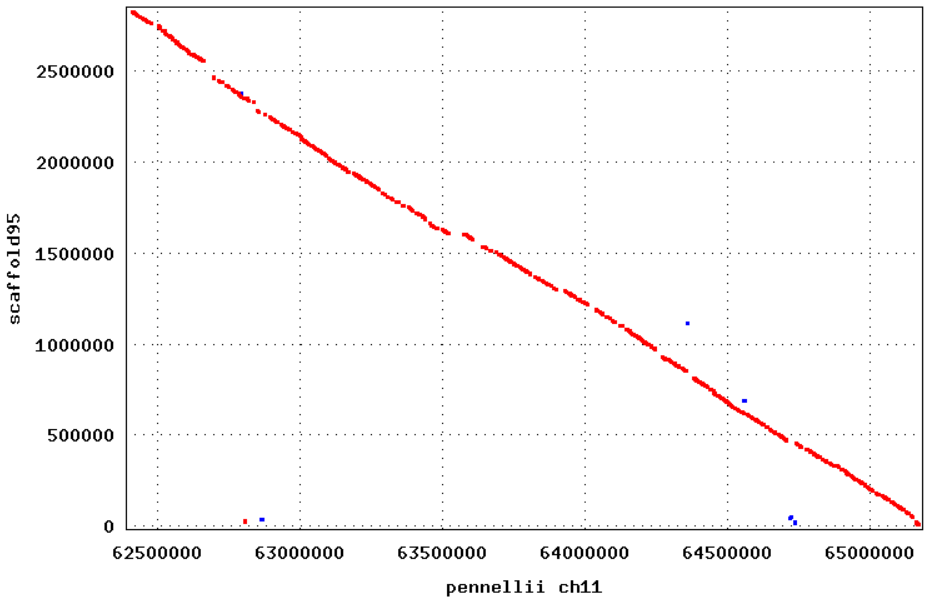

Supplementary Figure 6: Dotplots of the S. sitiens inversions against S. lycopersicum v3.0 and S pennelli, done with Mummer v4. The sequences were aligned with nucmer, using the “—mum” and “-c 500” options. The dotplots were generated with mummerplot using the “-large” option.

**A.**

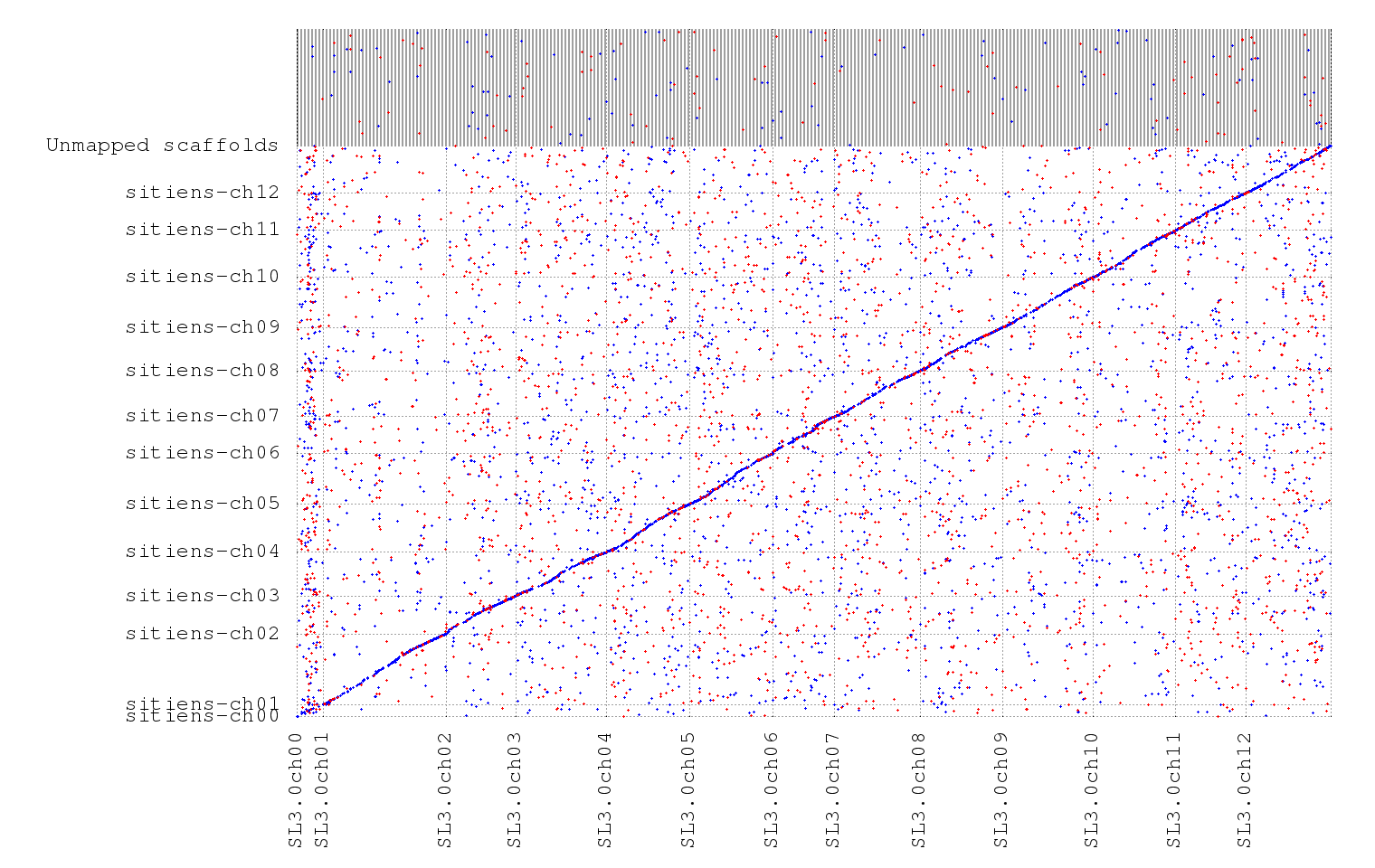

**B.**

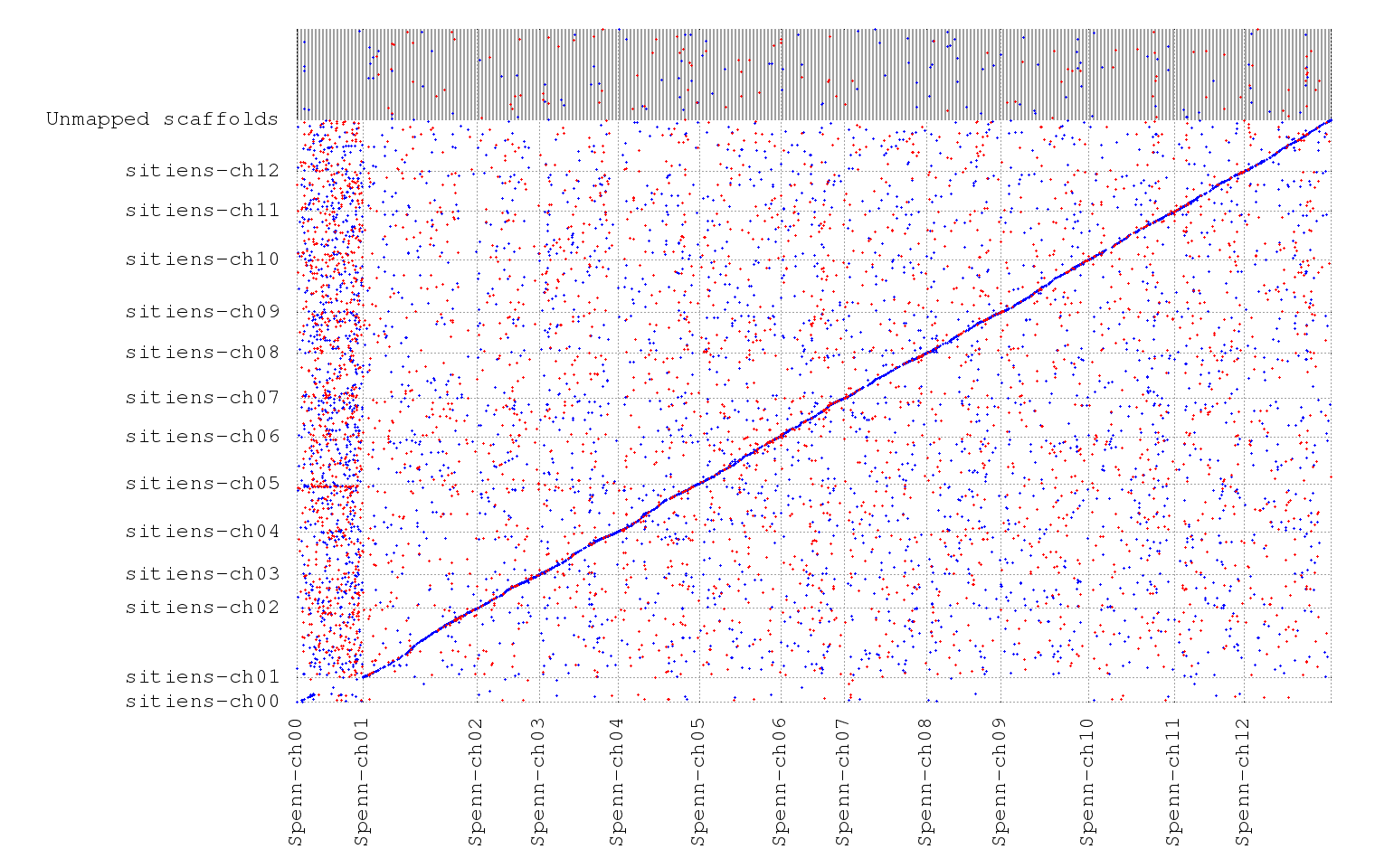

Supplementary Figure 7: Dotplots of the S. sitiens pseudomolecules assemblies against their respective references used in the chromosome_scaffolder script, S. lycopersicum v3.0 and S pennelli respectively. The assemblies were aligned with Mummer v4 using the “—mum” and “-c 500” options. The dotplots were generated with mummerplot using the “-large” option.

**1A.**
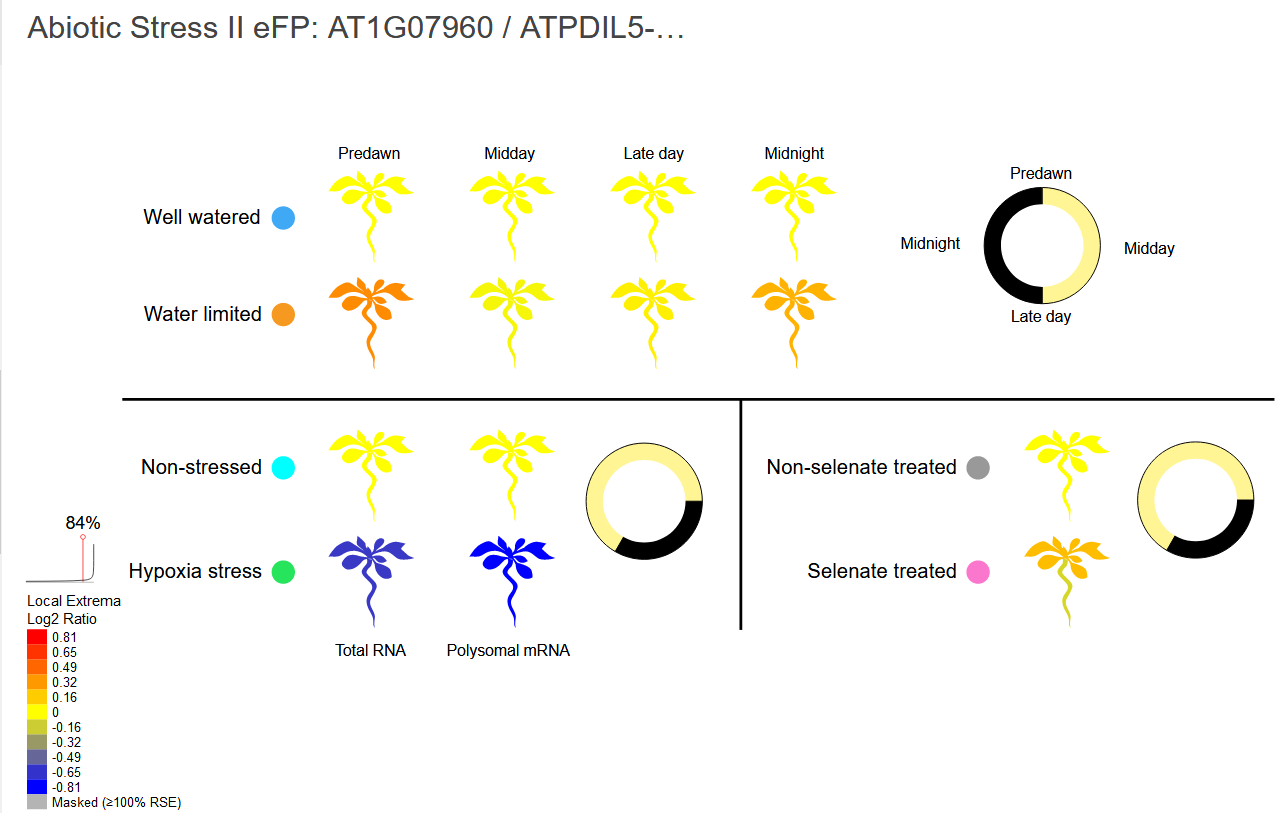

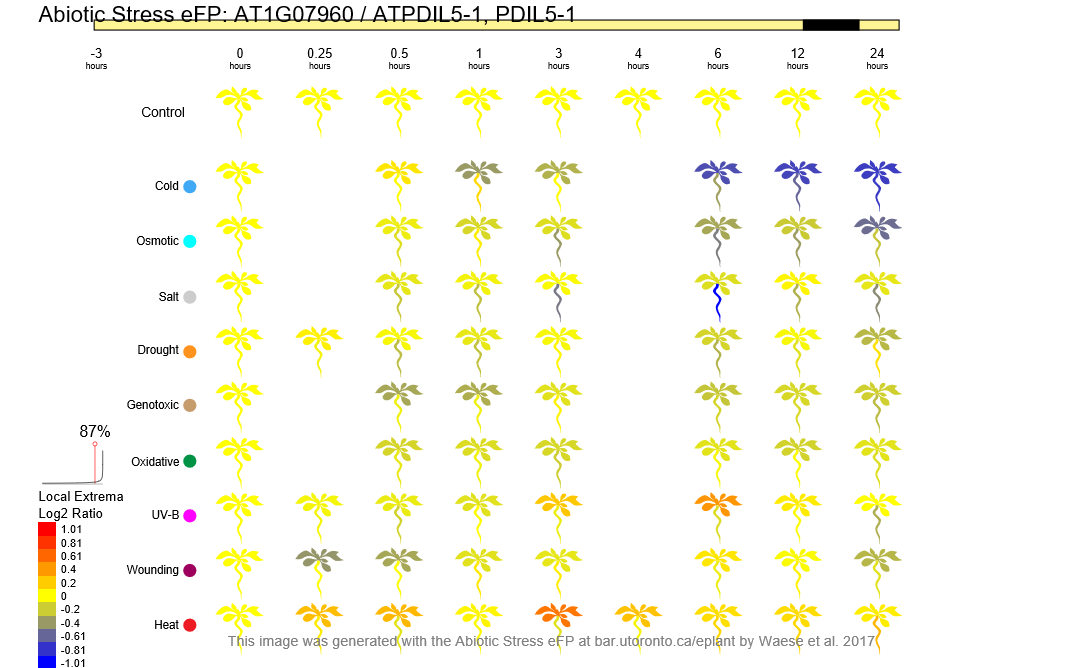
**1B.**

**2A.**

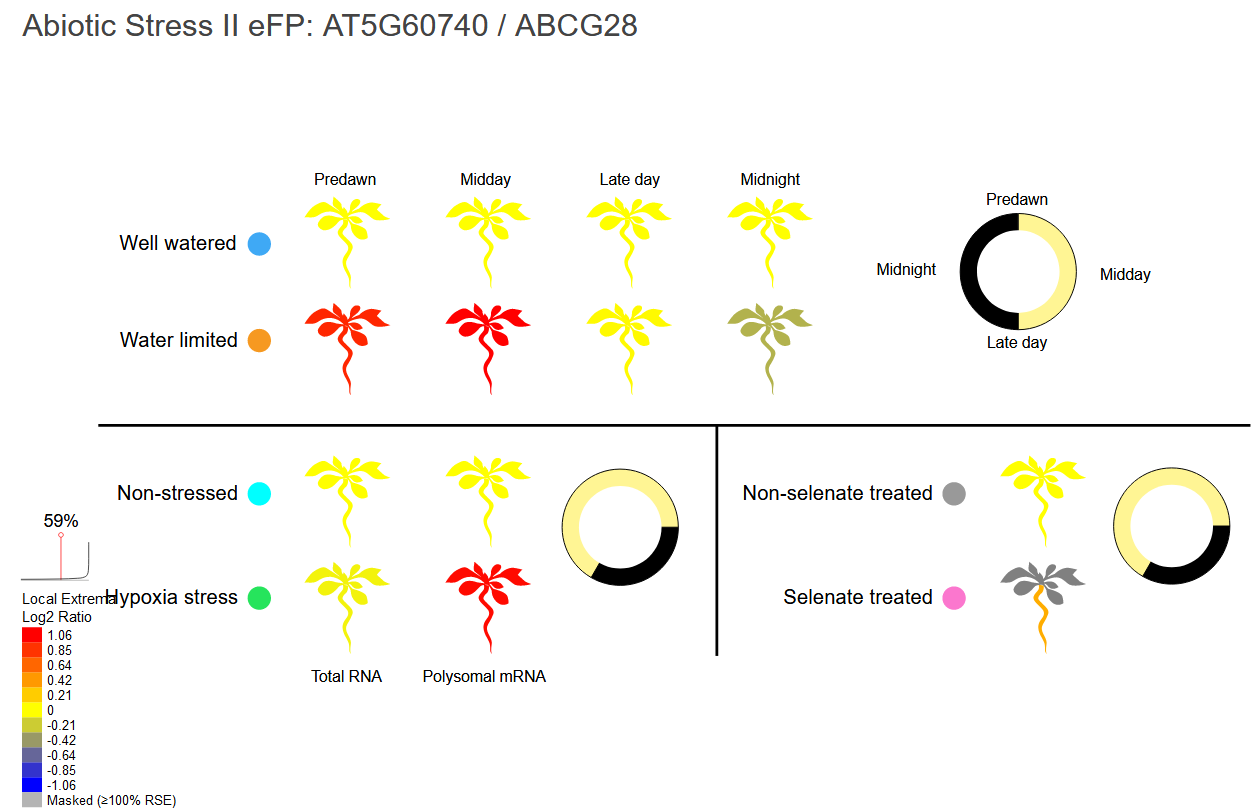

**2B.**

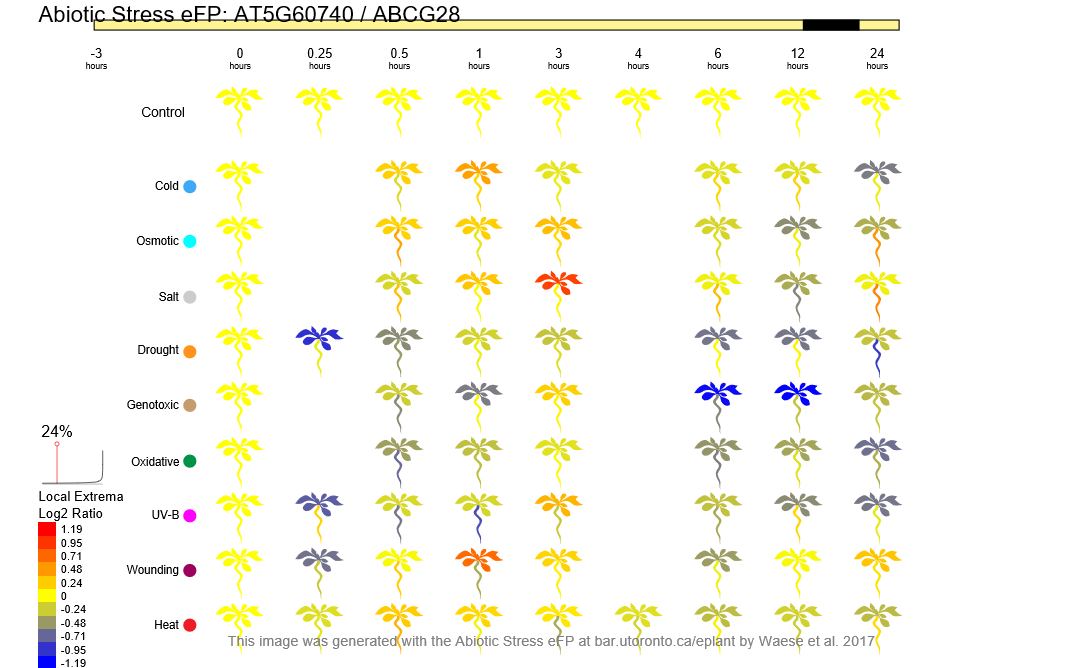

**3A.**

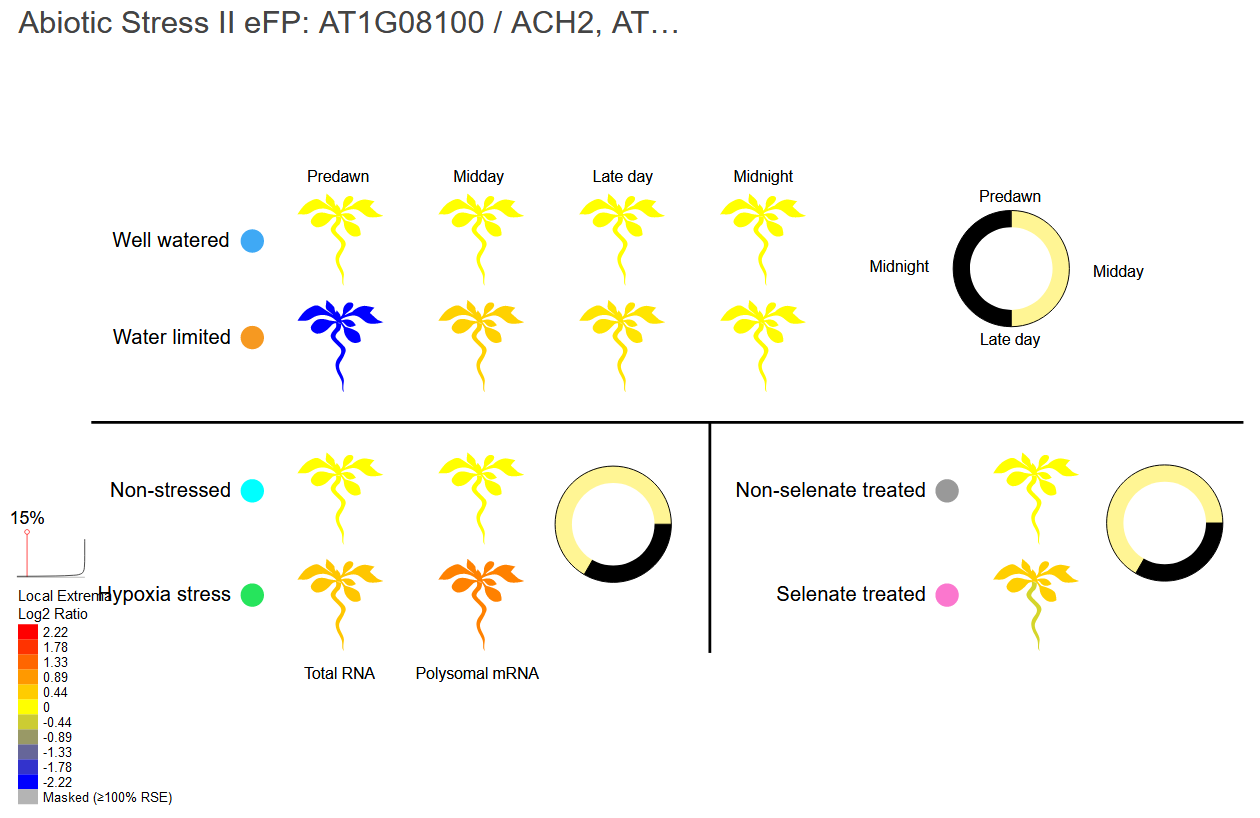

**3B.**
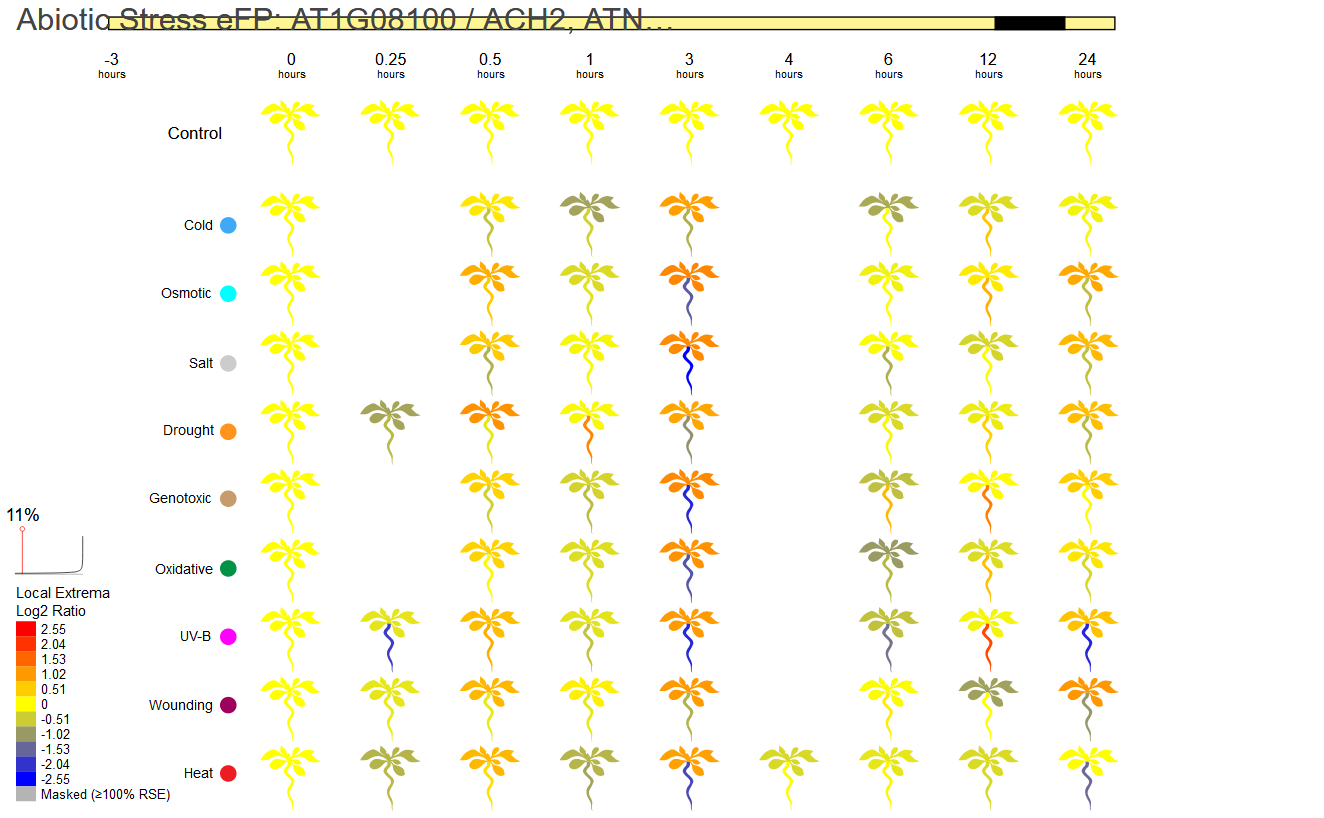

**4A.**

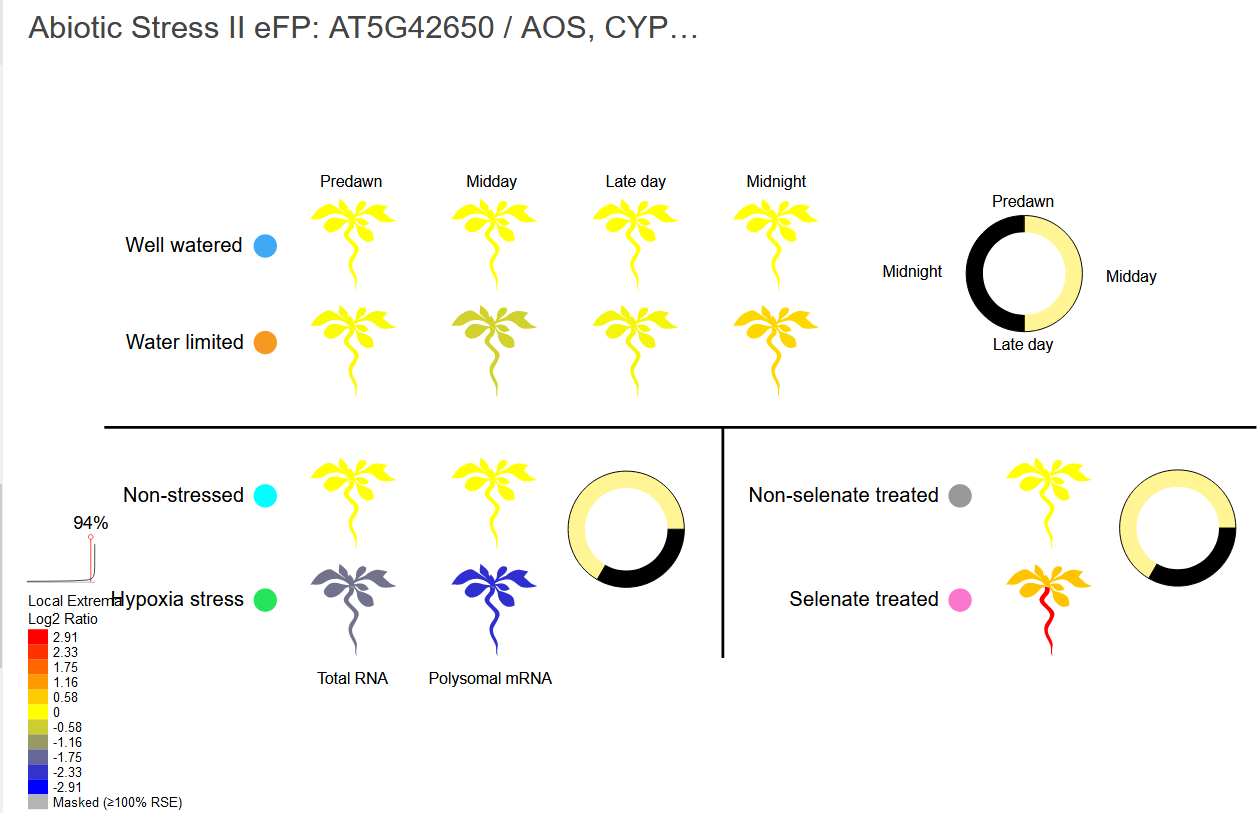

**4B.**

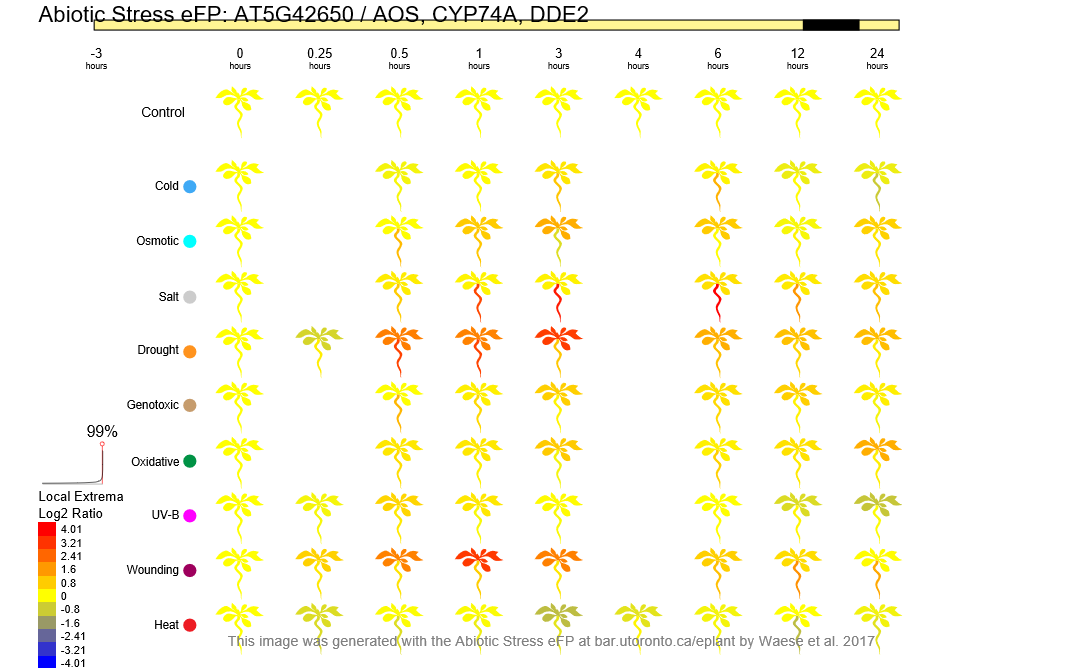

**5A.**

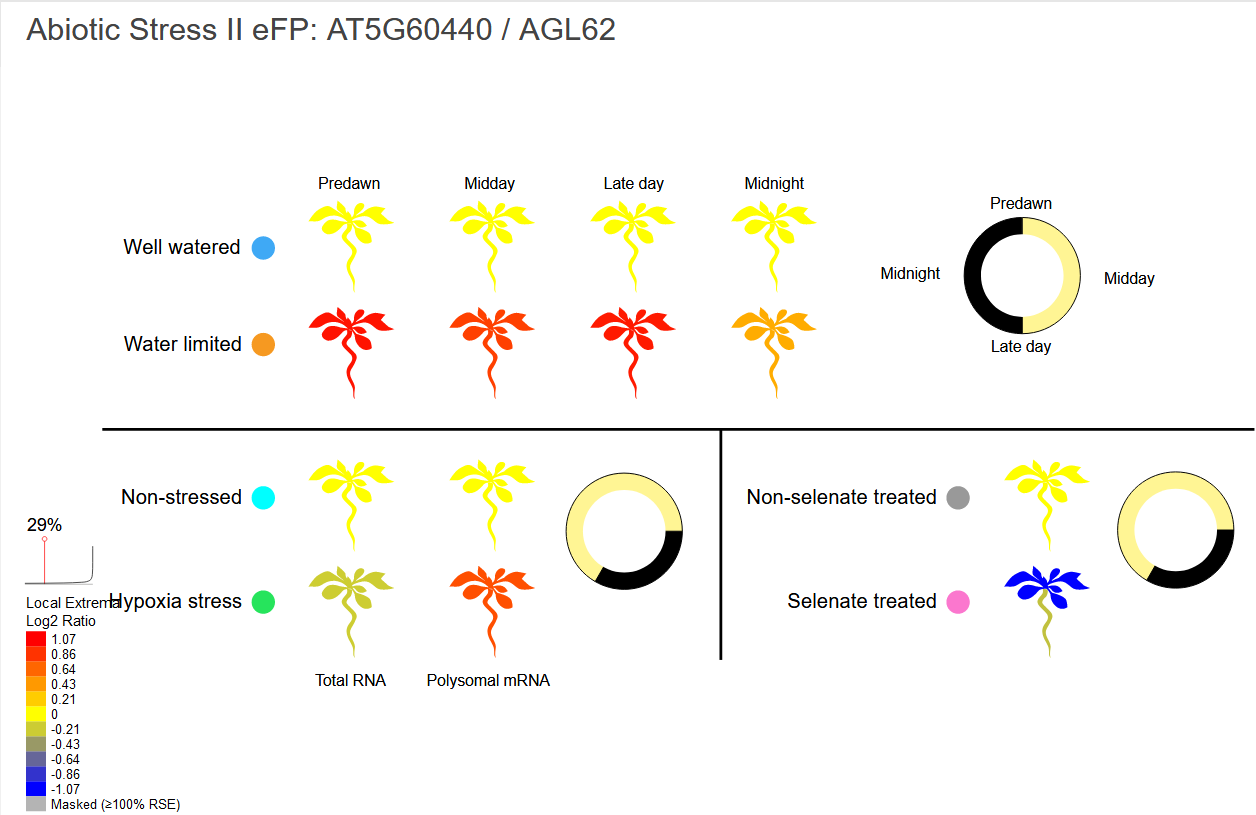

**5B.**

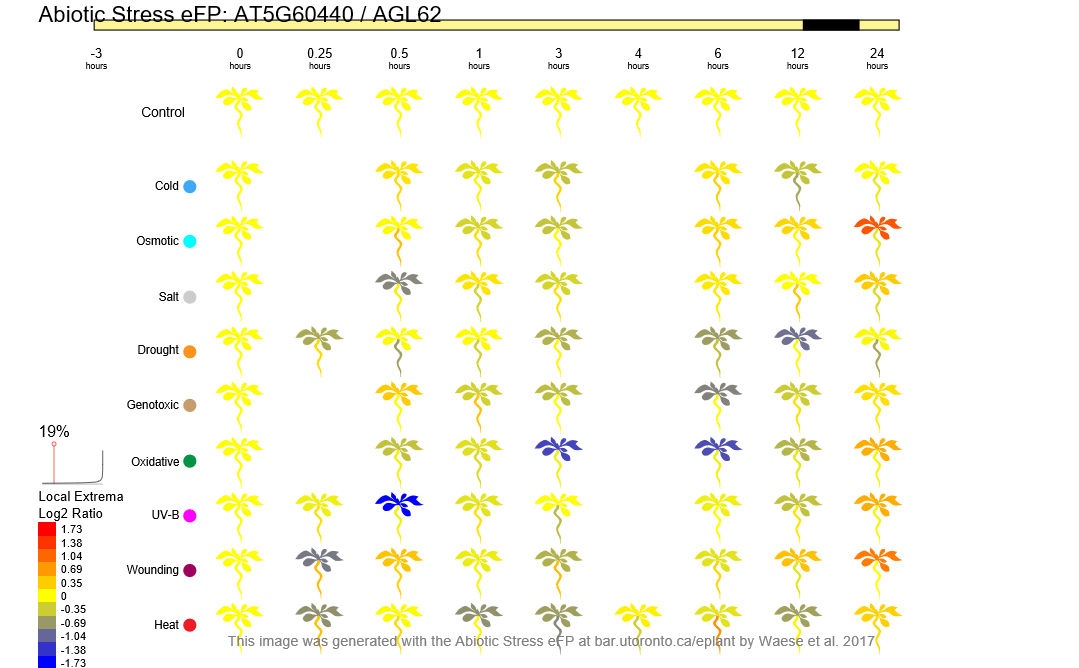

**6A.**

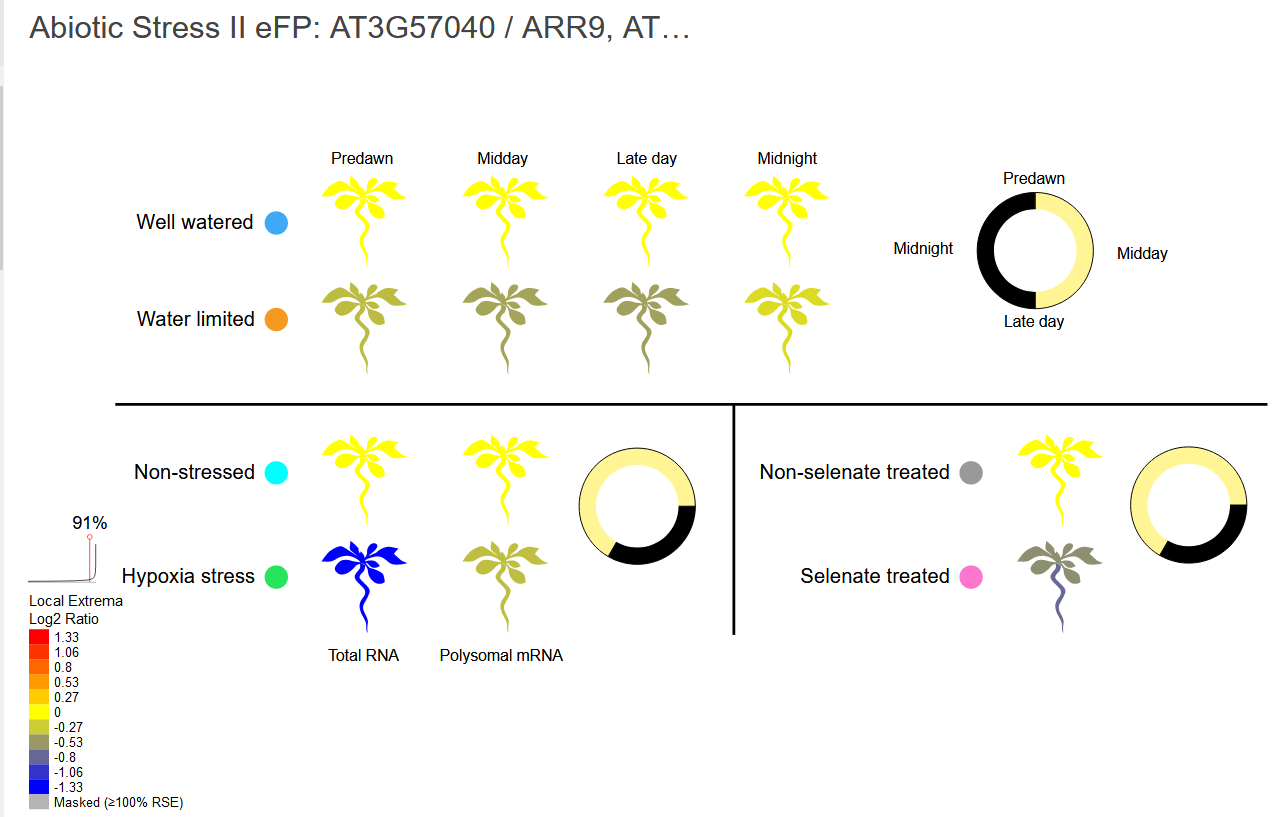

**6B.**

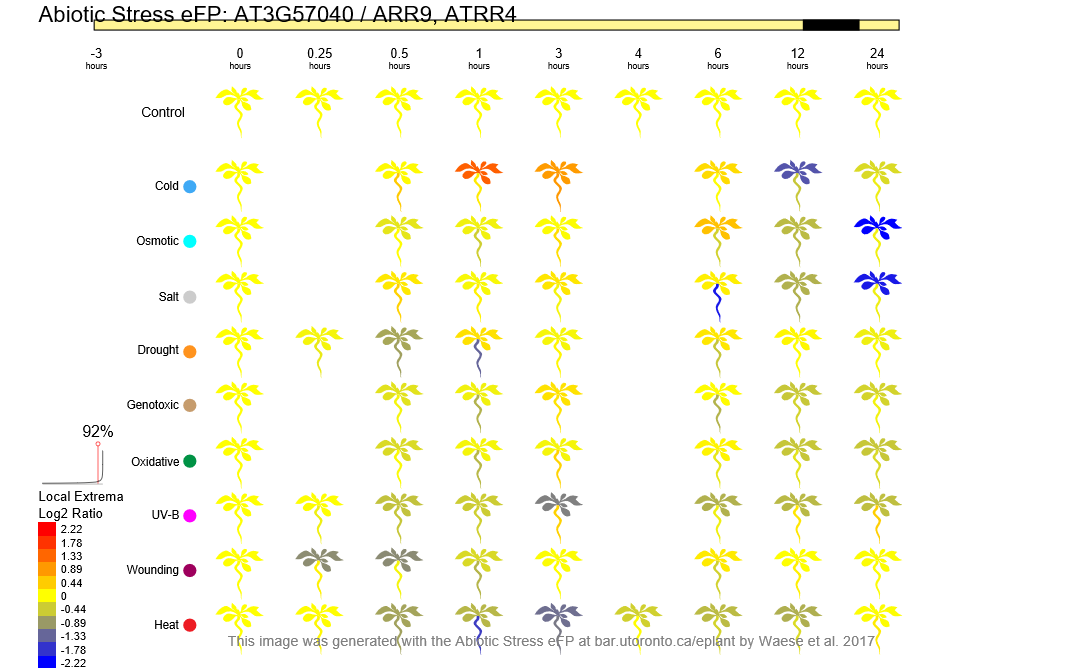

**7A.**

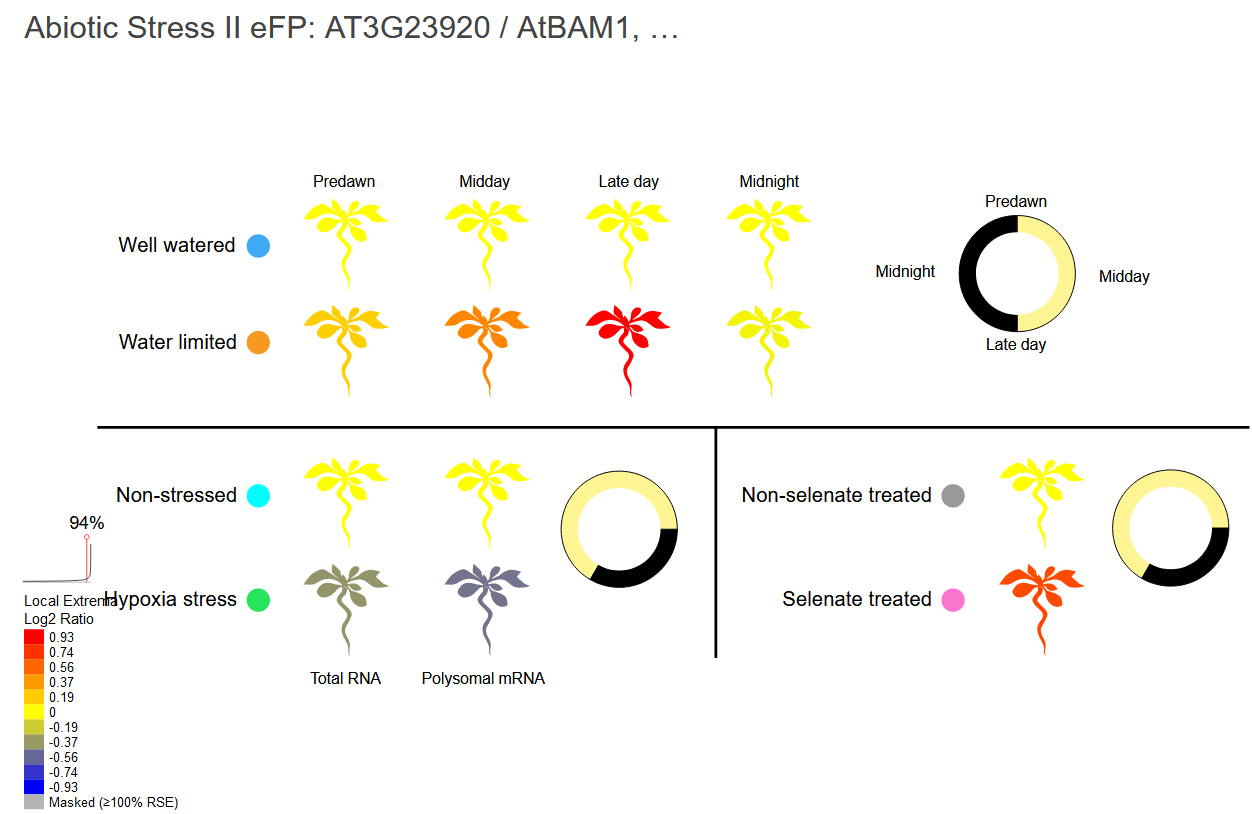

**7B.**

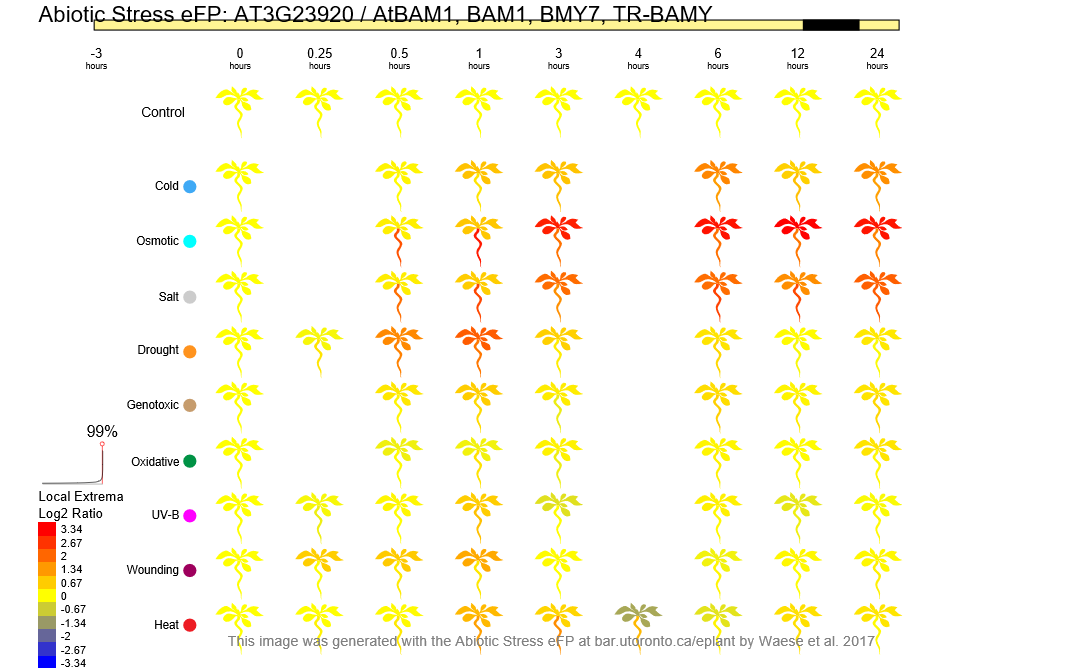

**8A.**

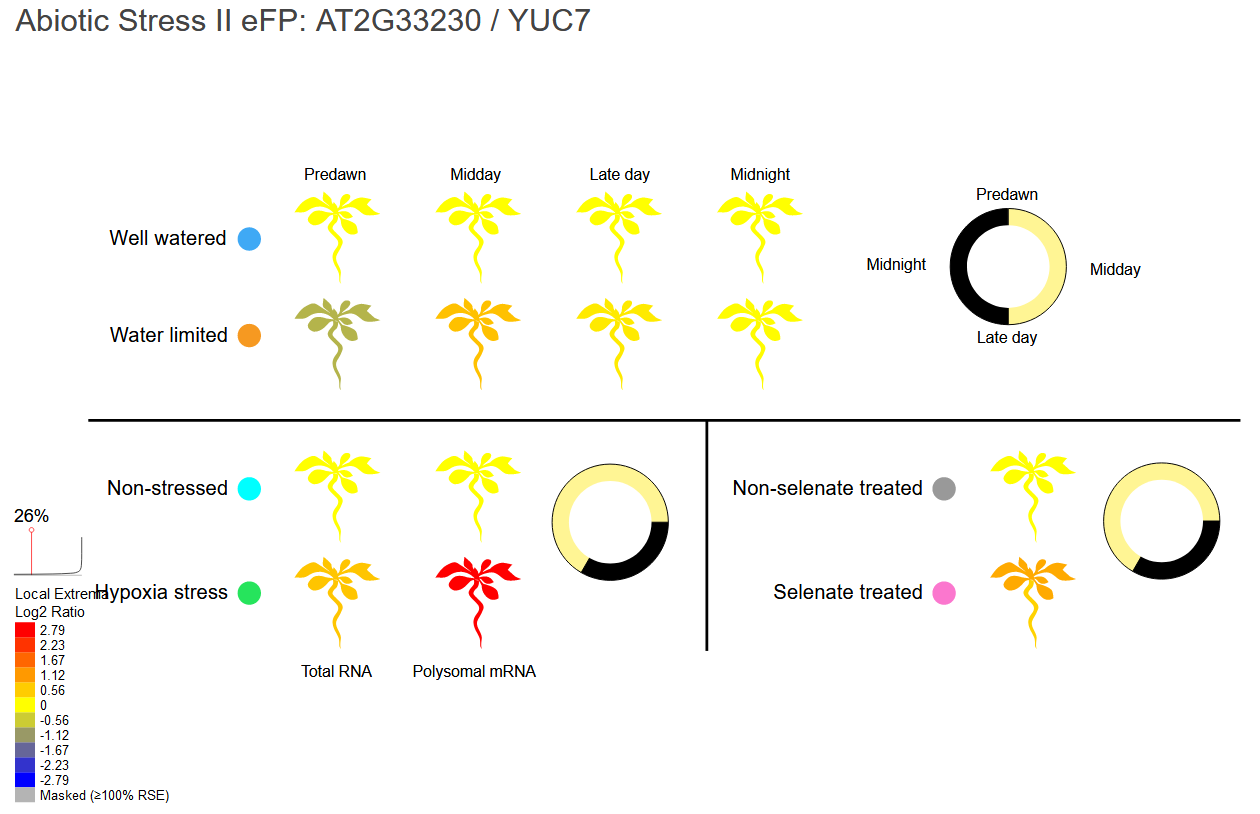

**8B.**

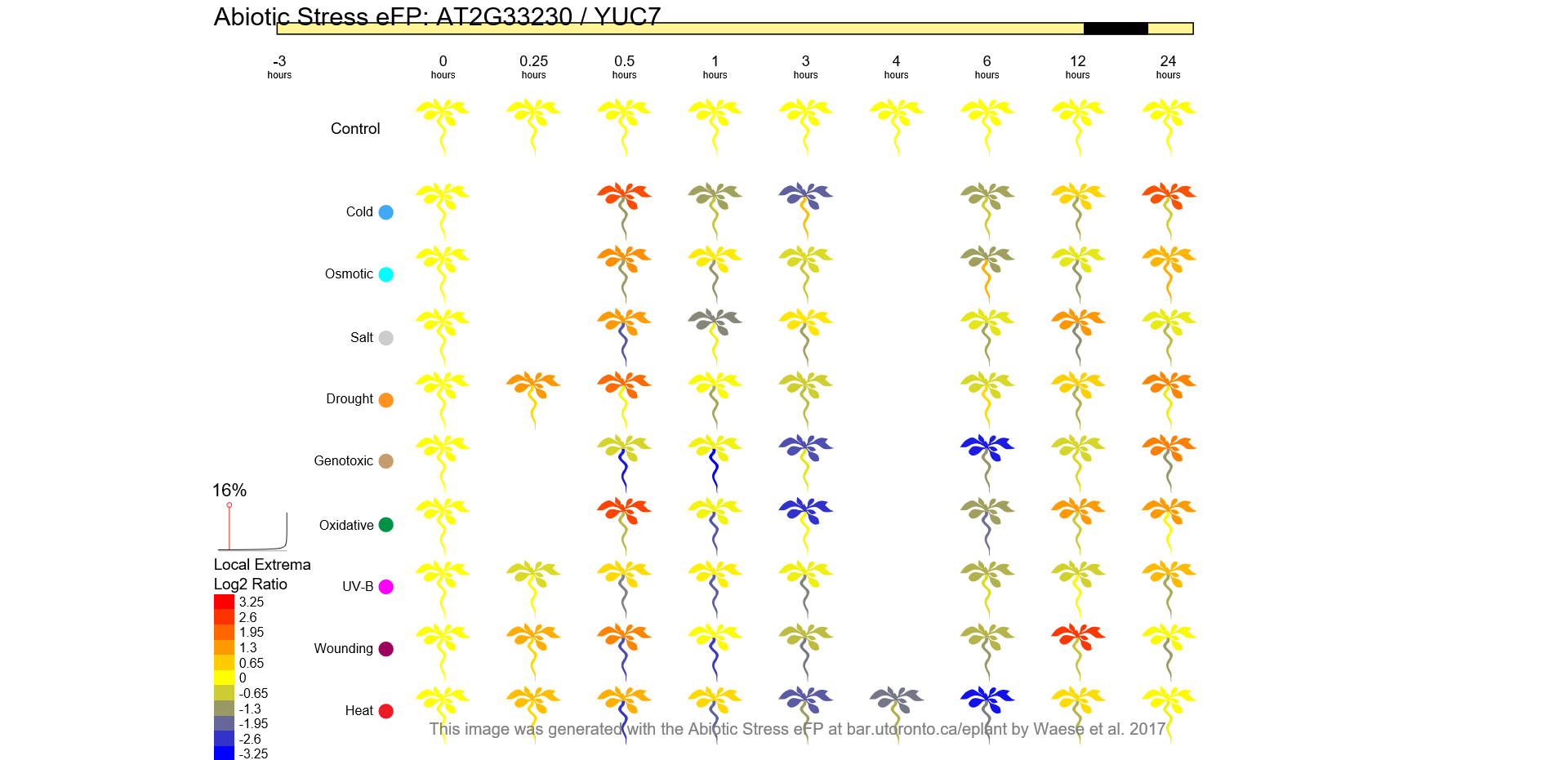

Supplementary Figure 8: Expression plots obtained from ePlant’s “Abiotic Stress II eFP” (**A**) and “Abiotic Stress eFP” (**B**) views under “Tissue & Experiment eFP viewers” for interesting genes identified in Solanum sitiens inversions against Solanum lycopersicum. Most of these genes are differentially expressed under salt and droughts stresses. **1.** AT1G07960 / Solyc11g069690 / Protein disulfide-isomerase 5-1 **2.** AT5G60740 / Solyc11g069820 / ABC transporter-like **3.** AT1G08100 / Solyc11g069735 / High-affinity nitrate transporter 2.2 **4.** AT5G42650 / Solyc11g069800 / Allene oxide synthase **5.** AT5G60440 / Solyc11g069770 / Agamous-like MADS-box protein AGL62 **6.** AT3G57040 / Solyc10g079600 / Two-component response regulator ARR9 **7.** AT3G23920 / Solyc09g091030 / beta-amylase 1, chloroplastic-like **8.** AT2G33230 / Solyc09g091090 / Probable indole-3-pyruvate monooxygenase YUCCA7
